# A multi-variant recall-by-genotype study of the metabolomic signature of body mass index

**DOI:** 10.1101/2021.10.21.465319

**Authors:** Si Fang, Kaitlin H. Wade, David A. Hughes, Sophie Fitzgibbon, Vikki Yip, Nicholas J. Timpson, Laura J. Corbin

## Abstract

**Objective:** We estimated the effect of body mass index (BMI) on circulating metabolites in young adults using a recall-by-genotype (RbG) study design.

**Methods:** An RbG study was implemented in the Avon Longitudinal Study of Parents and Children. Samples from 756 participants were selected for untargeted metabolomics analysis based on low/high genetic liability for higher BMI defined by a genetic score (GS). Regression analyses were performed to investigate the association between BMI GS groups and relative abundance of 973 metabolites.

**Results:** After correction for multiple testing, 29 metabolites were associated with BMI GS group. Bilirubin was amongst the most strongly associated metabolites with reduced levels measured in individuals with the highest BMI GS (beta=-0.32, 95% confidence interval (CI): -0.46, -0.18, Benjamini-Hochberg (BH) adjusted *p*=0.005). We observed associations between BMI GS group and levels of several potentially diet-related metabolites including hippurate which had lower mean abundance in individuals in the high BMI GS group (beta=-0.29, 95% CI: -0.44, -0.15, BH adjusted *p*=0.008).

**Conclusions:** Together with existing literature our results suggest a genetic predisposition to higher BMI captures differences in metabolism leading to adiposity gain. In the absence of prospective data, separating these effects from the downstream consequences of weight gain is challenging.

**Study Importance questions:** *What is already known about this subject?:*

- Metabolomics, defined as the measurement and study of circulating small molecules that are the substrates and products of cellular metabolism, is increasingly used by epidemiologists to provide a functional read-out of bulk cellular activity and a proxy to individual current health. This approach also provides insight into biological pathways linking exposures and disease.
- In observational studies, elevated body mass index (BMI) has been associated with a wide range of circulating metabolites. Researchers are now looking to genetic epidemiological methods, such as Mendelian randomization, to offer insight into potential causal relationships.

*What are the new findings in your manuscript?:*

- We identified 29 metabolites whose relative abundance varies with a genetic predisposition to higher BMI.
- Bilirubin, a key component of the heme catabolic pathway and a potent antioxidant, showed the strongest association with BMI score group.

*How might your results change the direction of research or the focus of clinical practice?:*

- Results of both Mendelian randomization and recall-by-genotype studies need to be combined with alternative study designs to distinguish between biomarkers that are intermediates on the pathway to BMI from those reflective of metabolic changes that result from increased adiposity.
- Separating causal biomarkers from non-causative biomarkers of adiposity is important since only the former are relevant to treatment and prevention, whilst both could be informative with respect to prediction and the downstream consequences of high BMI.

## Introduction

Despite the extensive focus in the literature, the full downstream impact of high body mass index (BMI) and the potential causal mechanisms by which BMI impacts a large number of non-communicable diseases remains unclear (1). Through a combination of large-scale observational studies and intervention designs, BMI has been established as a major risk factor for many common complex diseases, including type 2 diabetes mellitus, hypertension, myocardial infarction, stroke and several types of cancer (2–4). Whilst the precision of effect estimates describing the association between BMI and disease (and our confidence in them) has increased in line with greater sample sizes and independent replication, observational studies are limited by confounding, bias and reverse causation. Meanwhile, intervention studies (e.g., weight change protocols) designed to circumvent these conventional limitations have their own challenges – notably, a limited ability to alter BMI to the extent required to quantify an effect, and the necessarily short-term and small-scale nature of such interventions.

In response to these challenges and following developments in understanding genetic contributions to adiposity/BMI, methods from within the field of applied genetic epidemiology are now in use by researchers interested in dissecting the relationship between BMI and health. One such approach in which genetic variants act as an approximation to instrumental variables to evaluate the causal effect of an exposure (adiposity) on an outcome (disease), is Mendelian randomization (MR). In MR, genetic variation fulfils the role of an instrumental variable (5) where the presence of variance in BMI explained by genotype is (in principle) orthogonal to confounding factors and where genotype is assumed to exert an effect on health outcome only through BMI. Whilst validating the likely causal nature of the relationship observed between BMI and many common diseases, these studies do little to explain *how* the risk is delivered.

The study of circulating metabolites uses various techniques to detect and measure low molecular weight metabolites across a range of body fluids and tissues and can be used to provide a functional read-out of an individual’s current health. The use of metabolomics in epidemiological studies is increasing and has the potential to help elucidate the mechanisms linking obesity and associated co-morbidities, as well as identifying biomarkers to facilitate intervention and treatment. To date, studies have shown BMI-associated changes across a range of metabolite classes, including sex steroids, branched-chain and aromatic amino acids, acylcarnitines and lipids (6). But with much of the existing literature on the metabolomic impact of adiposity being based on observational epidemiological analyses, gaps remain in our understanding of the biology underpinning the development and direct pathophysiological consequences of obesity.

We aimed to integrate the use of genetic predictors for BMI with in-depth intermediate phenotyping to explore the relationship between BMI and metabolic health. Recall by genotype (RbG) is a study design in which participants (or samples) are selected from a pre-existing cohort based on genetic variation either at single variants or in the form of a genetic score (GS) (7). In this way, RbG exploits the concept of MR (i.e., the random assortment of genetic variants in offspring), enables greater power for a given number of samples analysed as compared to random selection, facilitates deep phenotyping and is typically less prone to confounding and reverse causality (7, 8). The aim of this RbG study was to examine the effect of BMI (the exposure) on circulating metabolites (the outcome) using a GS describing a high versus low predisposition for higher BMI.

## Methods

In this study, metabolomics data were derived from plasma samples collected during a routine research clinic conducted as part of the Avon Longitudinal Study of Parents and Children (ALSPAC). Individuals were selected for inclusion in the study based on a GS for BMI. The primary statistical analysis sought to identify metabolites whose levels were associated with GS group (low/high). Sensitivity analyses were performed to further characterise the associations observed. An overview of the study design is shown in **Figure 1**.

**Figure 1.**
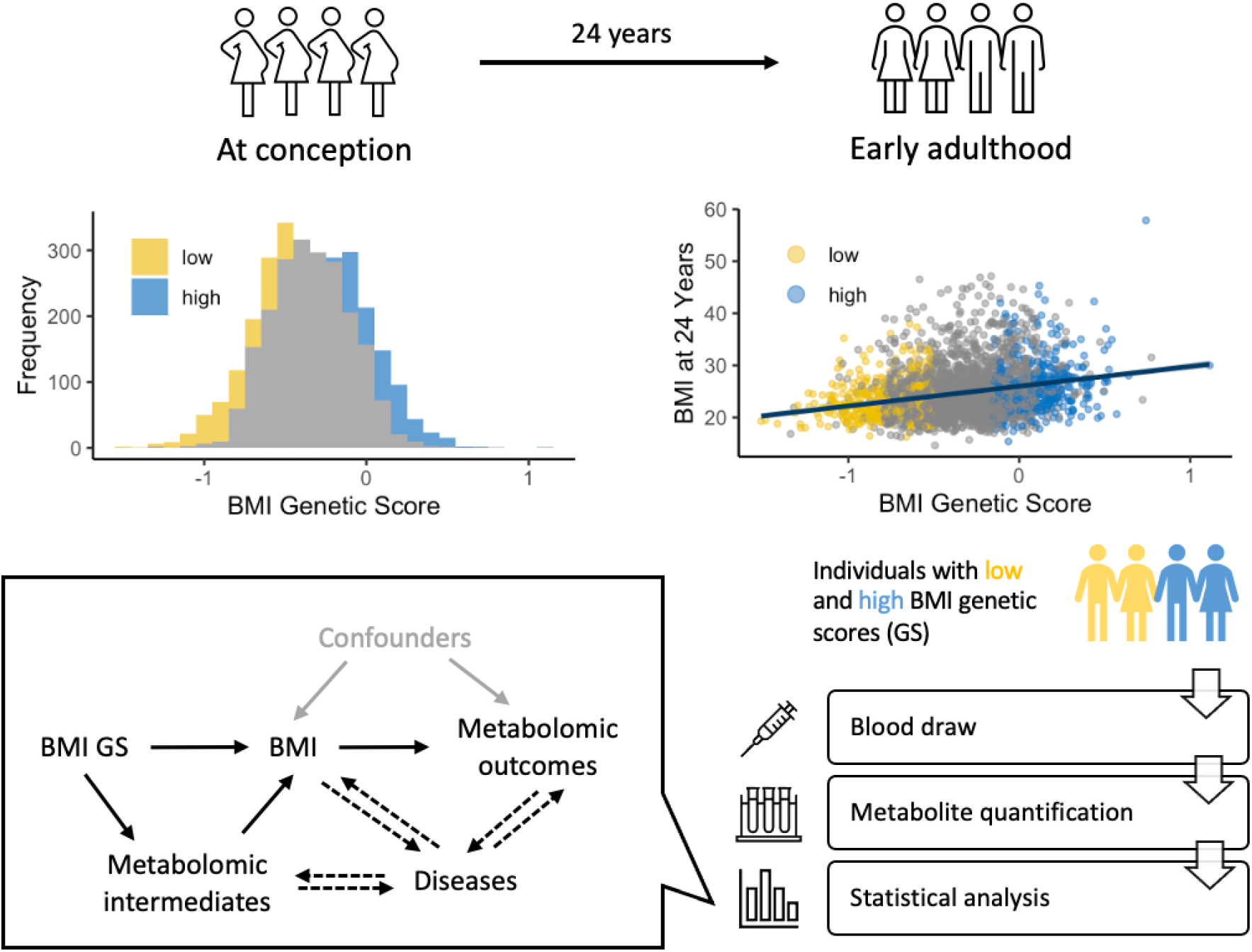
Study overview. This study involves the first-generation offspring in the Avon Longitudinal Study of Parents and Children (ALSPAC) multi-generational cohort, in which 14,541 pregnant women resident in the South West of England were recruited in the 90s. Firstly, we constructed genetic scores (GS) for body mass index (BMI) for all first-generation offspring. Under the recall-by-genotype study design, we recalled the plasma samples (collected at the age 24 years clinic) of individuals with a low (yellow) or high (blue) BMI GS for further analysis. Then, metabolites in those plasma samples were quantified by Metabolon. Finally, we performed statistical analysis to compare the metabolites levels between the two BMI GS groups. Our results are relevant to understanding the role of metabolites both as intermediates on the pathway to BMI and from BMI to disease.

### Study participants

ALSPAC is a prospective birth cohort of 14,541 pregnant women residing in the former region of Avon (UK) with expected dates of delivery from 1^st^ April 1991 to 31^st^ December 1992 (9–11) (see **Supplementary Methods** for a cohort summary). 13,988 children of the initial pregnancies who were alive at one year of age, referred to herein as Generation 1 (G1), have been followed up with a series of questionnaires and phenotypic assessments carried out during clinic visits. The study website contains details of all the data that is available through a fully searchable data dictionary and variable search tool (http://www.bristol.ac.uk/alspac/researchers/our-data/). For our analysis, plasma samples and phenotype data from a subset of G1 participants and selected phenotype data for their parents were included. Ethical approval for the study was obtained from the ALSPAC Ethics and Law Committee and the Local Research Ethics Committees (http://www.bristol.ac.uk/alspac/researchers/research-ethics/). Consent for biological samples was collected in accordance with the Human Tissue Act (2004). Informed consent for the use of data collected via questionnaires and clinics was obtained from participants following the recommendations of the ALSPAC Ethics and Law Committee at the time.

### Genotyping and sample selection

A subset of ALSPAC G1 participants (N=8,953) were genotyped using the Illumina HumanHap550 quad chip and data imputed to the 1000 Genomes reference panel (Phase 1, Version 3) (for full details see **Supplementary Methods**). A weighted GS was calculated for all G1 with genetic data using a previously published set of 940 near-independent genome-wide significant BMI-associated SNPs and their effect estimates (12). Following cross-matching against those G1 participants with data and samples collected at the age 24 years clinic visit, those with the highest and lowest GS were selected for inclusion in the study. In what follows, these GS-derived groups will be referred to as the ‘high BMI’ and ‘low BMI’ score groups, respectively. In total 760 samples were sent for analysis, split equally between the high BMI and low BMI score groups. For further details of the ‘GS derivation’ and ‘sample selection’ procedure see corresponding headings in the **Supplementary Methods**.

### Derivation of metabolite data

Fasted blood samples were collected at the age 24 years clinic visit from all ALSPAC G1 individuals who provided informed consent. For further details of blood sampling procedures see **Supplementary Methods**. Plasma samples were shipped on dry ice to Metabolon, Inc. (Durham, North Carolina, USA) for untargeted metabolomics analysis using established protocols. The Metabolon analysis consisted of four independent ultra-high-performance liquid chromatography-tandem mass spectrometry (UPLC-MS/MS) runs. Further details of the methodology can be found in **Supplementary Methods** and in published work (13, 14). Metabolite screening identified 1216 biochemicals, including 948 known and 268 unknown (at the time of analysis) compounds. Original scale data normalized in terms of raw area counts (as supplied by Metabolon) was used.

### Phenotype data collection

In ALSPAC, regular clinic visits of subsets of G1 were carried out from 4 months to 24 years old, including assessment of their basic anthropometric measures. For this study, data were extracted for several variables to characterise the GS-derived groups. To validate the performance of the GS, data were extracted from all available timepoints for BMI (kg/m^2^) (calculated as (weight (kg) / height squared (m^2^)). Data were also extracted for other measures of adiposity; this included weight (kg) and variables derived from a dual energy x-ray absorptiometry (DXA) scan of body composition, specifically, total body fat mass (kg) and total body lean mass (kg).

According to the theory of MR and analogous to the situation in a randomised controlled trial, creating groups based on a GS for BMI (as opposed to using BMI itself), should ensure those groups do not differ in any other respect, meaning any downstream analyses should be free from confounding. However, in the presence of (unmeasured) population structure, associations can be induced between GS (and therefore score group) and some of the traditional confounders of the relationship between BMI (as an exposure) and selected outcomes (15). Therefore, data were extracted for several phenotypic correlates of observed BMI to check for associations with score group and to evaluate the potential for them to act as confounders in the primary analysis. Full details of all phenotypic variables can be found in **Supplementary Methods**.

### Statistical analysis

An overview of the statistical analysis is provided in **Figure 2**. In all analyses, the low BMI score group was treated as the reference such that estimated effects represent the difference in the high BMI group relative to the low BMI group. All analyses were conducted in RStudio (16) using R v4.0.2 (17).

**Figure 2.**
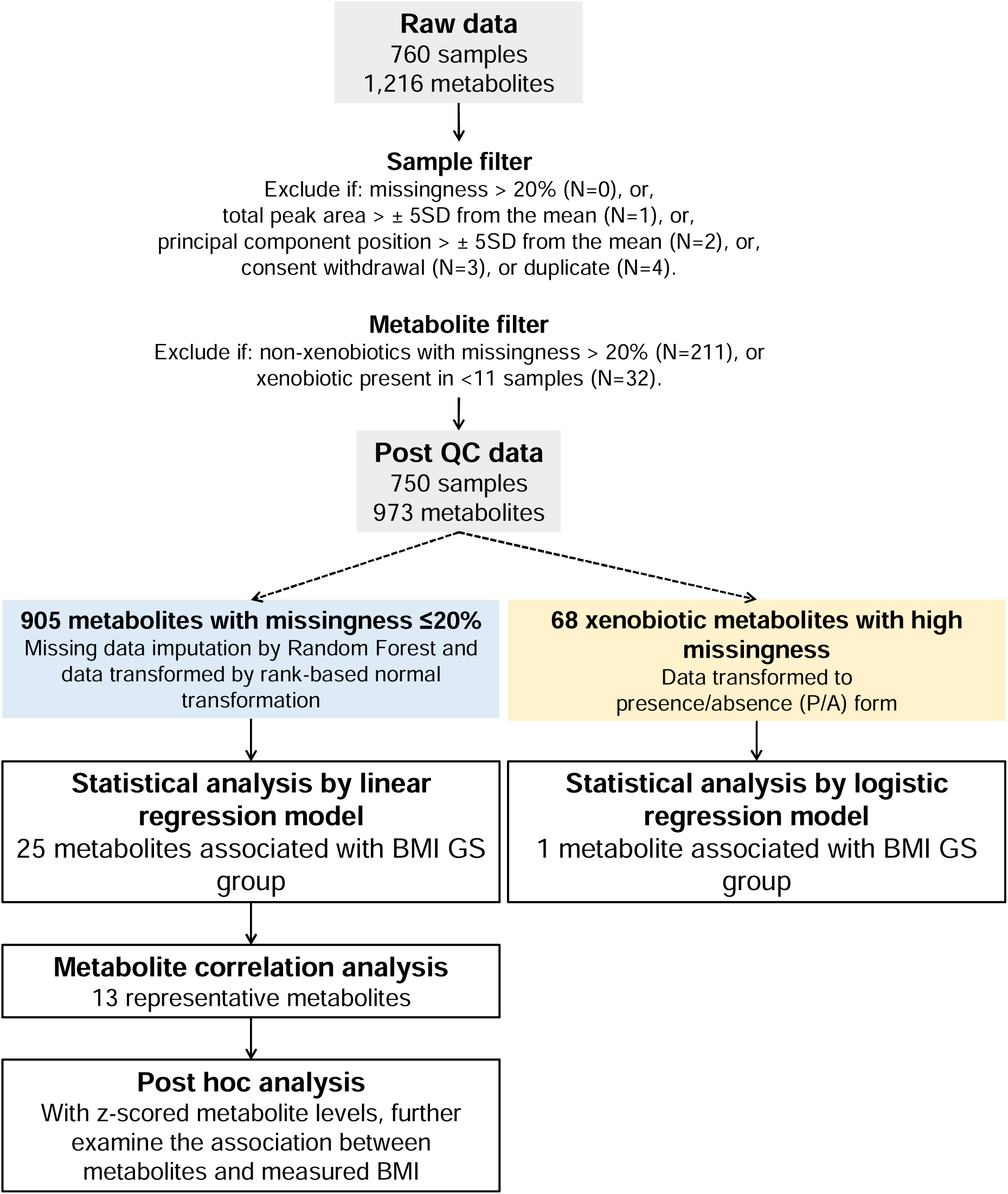
Overview of statistical analysis. ‘Raw data’ is the original scale data normalized in terms of raw area counts (as supplied by Metabolon). Data were prepared for statistical analysis by first filtering samples and metabolites based on a series of quality metrics and then applying imputation and re-scaling procedures as appropriate. SD = standard deviations; QC = quality control; BMI = body mass index; GS = genetic score.

#### Metabolite processing

We processed the raw (original scale) data received from Metabolon (N=760 samples) in preparation for statistical analysis using an in-house pipeline developed in R (17). Data were filtered based on a series of quality metrics (e.g., missing data). Except for those in the xenobiotic class (as annotated by Metabolon) metabolites with more than 20% missing values were excluded from the analysis. Following these exclusions, missing data were imputed using a random-forest based method and imputed data transformed using a rank-based normal transformation (RNT). In the case of xenobiotics (metabolites not produced by the human body), those with >20% missing were transformed to presence/absence (P/A) binary phenotypes such that missing values were replaced with 0 and recorded (non-zero) abundance measures replaced with 1. Xenobiotics were treated in this way because of their typically high level of missingness (or absence), which is both expected and biologically relevant given their (predominantly) exogenous origins. Xenobiotics present in <11 samples were excluded from downstream analyses on the basis that any statistical analyses would not be robust. A detailed description of this pre-analysis processing can be found in **Figure 2** and in **Supplementary Methods**. This procedure left 973 metabolites for analysis, including 905 continuous metabolites and 68 xenobiotics transformed to P/A.

#### Characterisation of recall groups

Between-group differences in our phenotype of interest, BMI (kg/m^2^), the previously described adiposity traits and potential confounders, including technical covariates, were assessed. The normal distribution of continuous variables was checked by a Shapiro-Wilk test and between-group differences assessed using a Student’s (two-sample, two-sided) t-test assuming unequal variance. In the case of BMI and weight, between-group differences were evaluated at different ages, ranging from four months to 24 years (when the samples used in this study were collected). Between-group differences in body composition measures were assessed based on DXA scans conducted during the age 24 years clinic. Between-group differences of continuous variables that could confound the primary analysis were also assessed by t-test. A Fisher’s exact test for count data was applied to test for possible association of group with binary and categorical variables.

#### Primary analysis - association of metabolites with recall group

To identify metabolite levels that differed between the low BMI and high BMI score groups, mean abundance was compared between groups using regression models. In Model 1, the post-imputation RNT metabolites were analysed within a linear regression framework [metabolite ∼ BMI.score.group]. The *R*^*2*^ from the model was used as an indication of the variance explained by score group. Log_2_ median fold change calculated as the ratio of median abundance (untransformed and unimputed) in the high BMI score group divided by median abundance in the low BMI score group was used to indicate relative effect sizes.

In Model 2, metabolites in the xenobiotic class with high levels of missingness and previously transformed to P/A traits, were analysed within a logistic regression framework [metabolite ∼ BMI.score.group]. In this case, the variance explained by the model was estimated using the ‘rsq’ function in the R package of the same name (18). A Benjamini-Hochberg (BH) correction was applied to adjust the p-values obtained from each of these analyses (Model 1 and Model 2) for multiple testing.

#### Sensitivity analyses

Several sensitivity analyses were carried out to aid interpretation and to further characterise the associations observed in the primary analysis. These analyses were restricted to the subset of BMI score group associated metabolites (BH p<0.05) output from Model 1. A full description of the methods used can be found under corresponding headings in **Supplementary Methods**.

##### Extension of primary association analyses

Model 1 was extended to a multivariate model in which any potential confounder that had previously been shown to be associated with score group was fitted as an independent fixed effect alongside score group.

##### Metabolite correlation analysis

A hierarchical clustering approach was applied to the subset of associated metabolites to identify redundancy in the data (i.e., where associated metabolites were highly correlated and likely representing the same biological signal). A reduced set of ‘representative’ metabolites was derived forming the focus for the next steps.

##### Association of metabolites with measured BMI

Linear regression analyses were conducted to evaluate the direct association between measured BMI (at the age 24 years clinic) and the subset of BMI score group associated metabolites with BMI score group, sex and age fitted as covariates in a multivariate linear model [metabolite ∼ BMI + BMI.score.group + sex + age]. In order to investigate the consistency of the BMI effect across the two groups, the same model was also fitted with an interaction term [metabolite ∼ BMI * BMI.score.group + sex + age] and within each BMI score group separately [metabolite ∼ BMI + sex + age].

## Results

After filtering based on a series of pre-defined quality metrics, the study sample consisted of samples from 750 G1 individuals with abundance measures for 973 metabolite traits (905 continuous and 68 presence/absence traits). Details of the exclusions made are shown in **Figure 2** and described in **Supplementary Methods (‘Metabolite data processing’)**. The phenotypic characteristics of all G1 individuals who attended the age 24 years clinic as compared to the study sample (after QC) are presented in **Table 1**. Both recall groups were consistent with the overall cohort in terms of age and sex distribution, whilst adiposity traits showed expected differences (see below for details).

**Table 1.** Characteristics of participants based on data collected at the age 24-year clinic.

| | | All attending<sup>a</sup> (N=4019) | | Low BMI group (N=373) | | High BMI group (N=377) | | Between-group difference in proportion (OR)/mean [95% CI], $p^b$ |
| --- | --- | --- | --- | --- | --- | --- | --- | --- |
|  |  | Mean/n | St. Dev./% | Mean/n | St. Dev./% | Mean/n | St. Dev./% |  |
| **Sex** | **Male** | 1505 | 37.4% | 148 | 39.7% | 150 | 39.8% | 1.00 [0.74,1.36], $p=1.00$ |
|  | **Female** | 2514 | 62.6% | 225 | 60.3% | 227 | 60.2% |  |
| **Age (years)** | **Male** | 24.5 (n=1505) | 0.80 | 24.6 (n=148) | 0.80 | 24.5 (n=150) | 0.73 | -0.05 [-5.85,5.76]<sup>c</sup>, $p=0.99$ |
|  | **Female** | 24.5 (n=2514) | 0.82 | 24.4 (n=225) | 0.75 | 24.4 (n=227) | 0.81 |  |
| **BMI (kg/m<sup>2</sup>)** | **Male** | 24.9 (n=1460) | 4.44 | 23.8 (n=148) | 3.46 | 26.2 (n=150) | 4.66 | 2.78 [2.12,3.43], $p=3.79 \times 10^{-16}$ |
|  | **Female** | 25.0 (n=2479) | 5.42 | 23.1 (n=222) | 3.79 | 26.1 (n=223) | 5.62 |  |
| **Weight (kg)** | **Male** | 80.6 (n=1495) | 15.3 | 78.5 (n=148) | 12.6 | 85.3 (n=150) | 17.1 | 8.01 [5.72,10.3], $p=1.56 \times 10^{-11}$ |
|  | **Female** | 68.8 (n=2481) | 15.8 | 63.4 (n=223) | 10.8 | 72.1 (n=223) | 16.4 |  |
| **Total fat mass (kg)** | **Male** | 20.6 (n=1459) | 9.77 | 18.6 (n=146) | 7.94 | 23.4 (n=144) | 10.8 | 5.67 [4.26,7.07], $p=1.17 \times 10^{-14}$ |
|  | **Female** | 25.1 (n=2403) | 11.1 | 21.0 (n=217) | 7.79 | 27.3 (n=212) | 10.9 |  |
| **Total lean mass (kg)** | **Male** | 56.9 (n=1459) | 7.55 | 57.2 (n=146) | 7.05 | 59.1 (n=144) | 8.20 | 2.12 [0.60,3.63], $p=6.33 \times 10^{-03}$ |
|  | **Female** | 41.2 (n=2403) | 5.41 | 40.0 (n=217) | 4.46 | 42.1 (n=212) | 5.04 |  |

BMI = body mass index; OR = odds ratio; <sup>a</sup> Summary statistics based on all those who attended the age 24 years clinic; <sup>b</sup> results from a Student's two-sample two-sided t-test to compare means in the high BMI group as compared to the low BMI group. In the case of sex, a Fisher's test was performed to test for a difference in the proportion of males versus females in the two groups and the results presented as an odds ratio; <sup>c</sup> the test for a difference in mean age was conducted based on age in weeks at the age 24 years clinic.

### Characterisation of recall groups

At the age 24 years clinic when the samples were collected, the mean (standard deviation, SD) BMI of individuals in the low BMI score group was 23.4 (3.7) kg/m^2^, falling within the ‘normal weight’ range as defined by the World Health Organisation (18.5kg/m^2^ to <25kg/m^2^). In contrast, the mean BMI of individuals in the high BMI score group, 26.1 (5.2) kg/m^2^, fell within the ‘pre-obesity’ range (25kg/m^2^ to <30kg/m^2^). Differences were also observed in weight and in both total fat mass and total lean mass at this timepoint (**Table 1**). Temporal analyses showed that the between-group differences in BMI emerged at about four years of age then increased rapidly until participants reached around 13 years of age and somewhat more slowly thereafter (**Figure 3** and **Supplementary Table S1**); a similar pattern was observed in weight (**Figure S1** and **Supplementary Table S1**). There was little evidence for an association between BMI score group and most potential confounders tested (**Supplementary Table S2**). BMI score group showed modest association with parental (mothers and mother’s partners) social class (**Supplementary Table S2**) but with no clear direction of effect across categories (**Supplementary Figure S2**).

**Figure 3.**
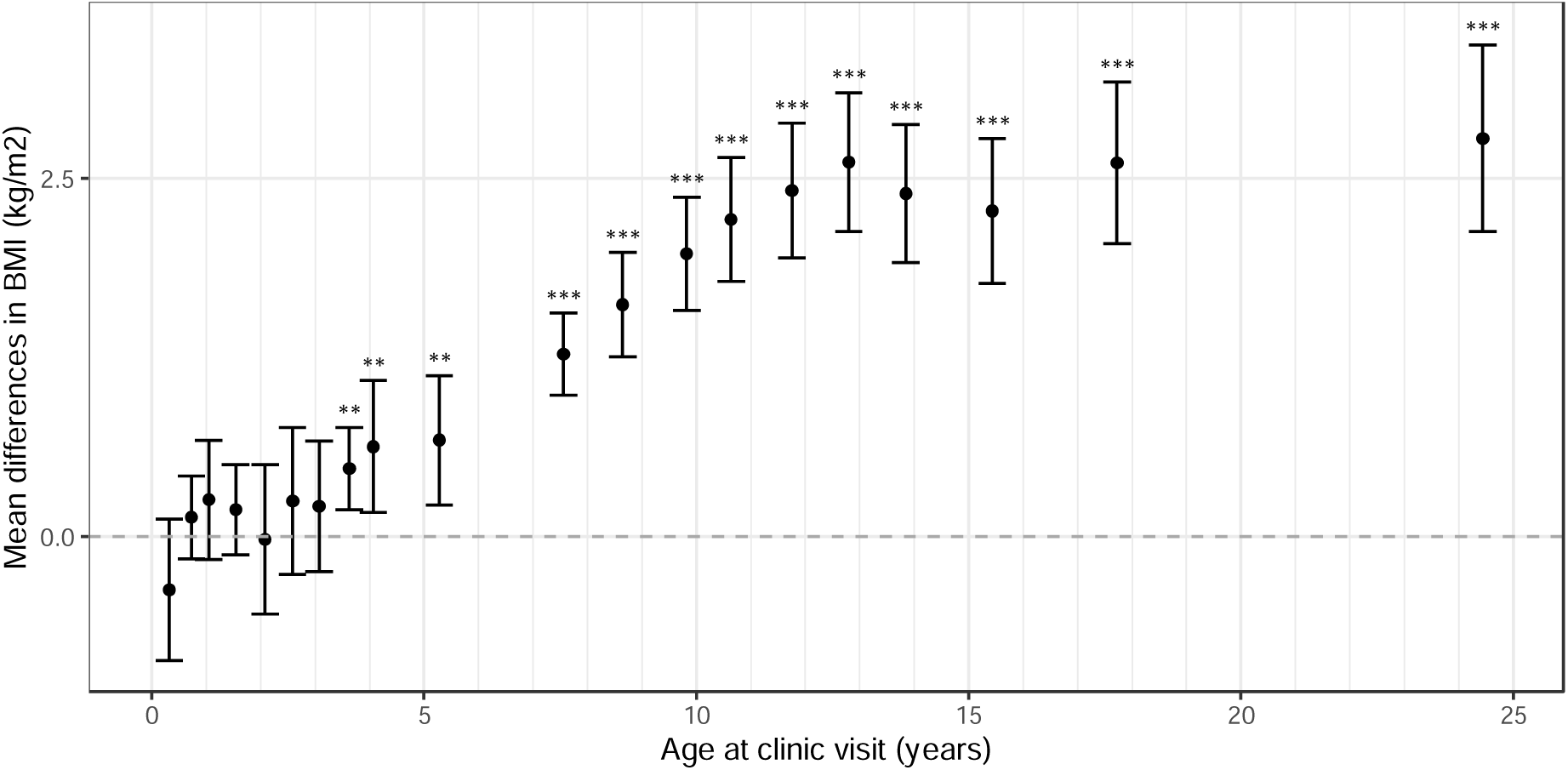
Mean differences in BMI between the high and low BMI score groups. Error bars represent the standard errors of mean differences in weight. Sample size ranges from 108 (at age 31 months) to 743 (at age 24 years). Test results are given for a Students (two-sample, two-sided) t-test. ***: p-value < 0.001; **: p-value < 0.01; *: p-value < 0. For full results see **Supplementary Table S1**.

### Association of metabolites with recall group

Overall, we observed relatively small differences across a wide range of molecules with median log_2_ fold changes typically in the range -0.5 to 0.5 and a slight bias towards decreased abundances in the high BMI score group (**Figure 4**). Of the 905 metabolites tested in Model 1, 29 were associated with BMI score group (BH *p*<0.05), 25 of which had annotations available from Metabolon (as of February 2020) (**Table 2**) (see **Supplementary Table S3** for full results). 25 of 29 (86%) had lower mean abundance in the high BMI score group compared to the low BMI group. The four metabolites that show the greatest evidence for association with BMI score group were bilirubin and bilirubin degradation products from the “Haemoglobin and Porphyrin Metabolism” pathway. A total of 11 metabolites assigned to this pathway appeared in the list of associated metabolites, including biliverdin. Score group allocation explained 2.6% of the variation in the abundance of the most strongly associated bilirubin degradation product. Four metabolites showed a positive association with high BMI score group, including two forms of sphingomyelin and metabolonic lactone sulfate. For the 29 associated metabolites, within-group distributions of metabolite levels were visualised using box and whisker plots with the original (unimputed abundance) data after mean centring and scaling as input (**Supplementary Figure S3**).

**Figure 4.**
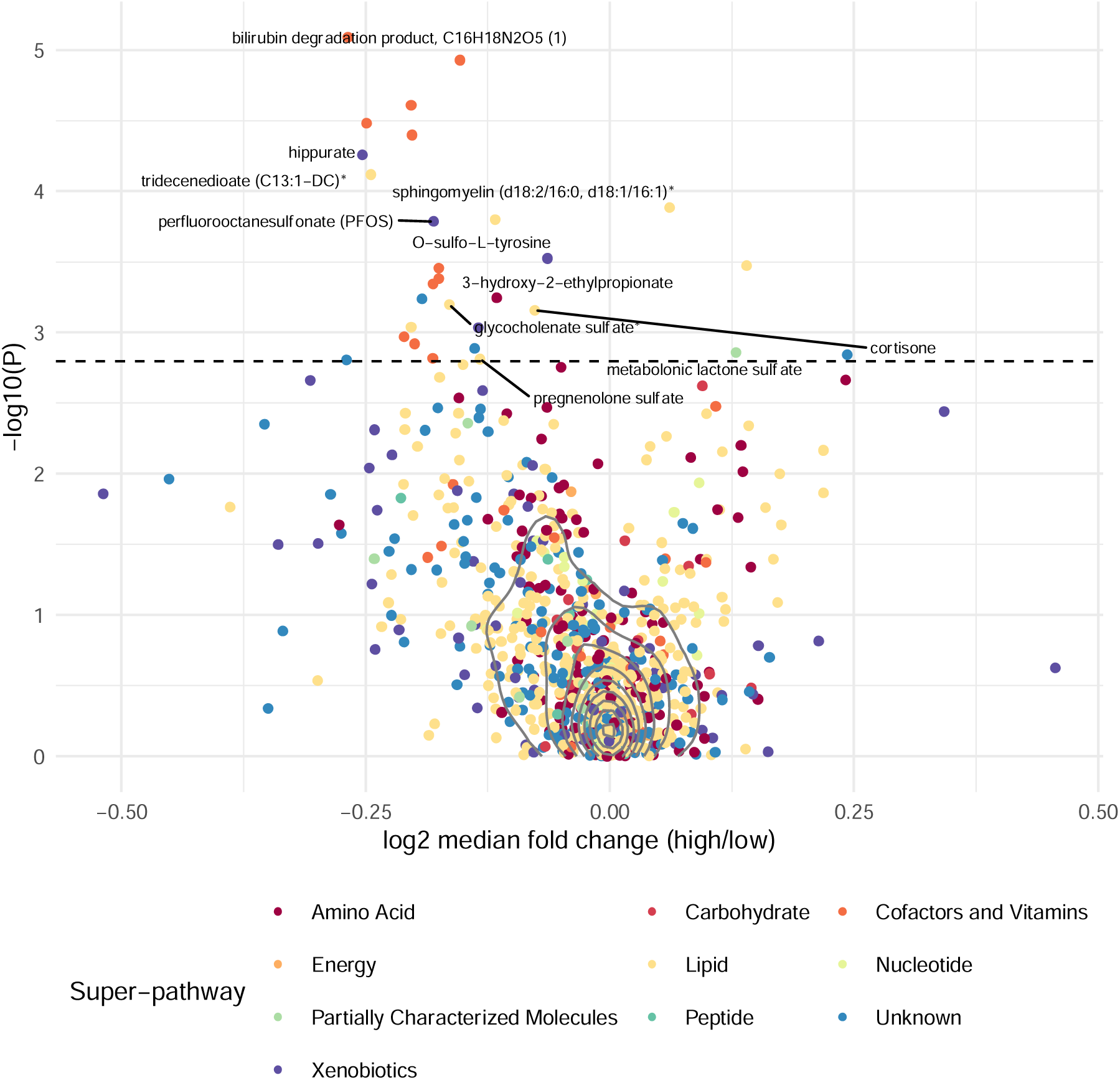
Volcano plot depicting the association between circulating metabolites and BMI score group. Points are coloured by super-pathway. Log_2_ median fold change calculated as the ratio of median abundance (untransformed and unimputed) in the high BMI score group divided by median abundance in the low BMI score group. P-values used to derive -log_10_(p) are those from the linear regression analysis. All points above the dashed line have a Benjamini-Hochberg adjusted p-value <0.05. Solid grey lines indicate the density of points. A representative selection of metabolites of known identity are labelled. *: Indicates a compound that has not been confirmed based on a standard.

**Table 2.** List of identified metabolites associated with BMI score group.

| Metabolite | Subpathway | Beta (95% CI) | BH *p* | Cluster<sup>a</sup> |
| --- | --- | --- | --- | --- |
| bilirubin degradation product, C16H18N2O5 (1) | Hemoglobin and Porphyrin Metabolism | -0.32 (-0.47,-0.18) | 0.005 | 1** |
| bilirubin (Z,Z) | Hemoglobin and Porphyrin Metabolism | -0.32 (-0.46,-0.18) | 0.005 | 1 |
| bilirubin (E,Z or Z,E)* | Hemoglobin and Porphyrin Metabolism | -0.31 (-0.45,-0.17) | 0.007 | 1 |
| bilirubin degradation product, C16H18N2O5 (2) | Hemoglobin and Porphyrin Metabolism | -0.30 (-0.44,-0.16) | 0.007 | 1 |
| biliverdin | Hemoglobin and Porphyrin Metabolism | -0.30 (-0.44,-0.16) | 0.007 | 1 |
| hippurate | Benzoate Metabolism | -0.29 (-0.44,-0.15) | 0.008 | 2 |
| tridecenedioate (C13:1-DC)* | Fatty Acid, Dicarboxylate | -0.29 (-0.43,-0.15) | 0.010 | 3** |
| sphingomyelin (d18:2/16:0, d18:1/16:1)* | Sphingomyelins | 0.28 (0.14, 0.42) | 0.015 | 4** |
| 3-decenoylcarnitine | Fatty Acid Metabolism (Acyl Carnitine, Monounsaturated) | -0.28 (-0.42,-0.13) | 0.015 | 3 |
| perfluorooctane sulfonate (PFOS) | Chemical | -0.27 (-0.42,-0.13) | 0.015 | 5 |
| O-sulfo-L-tyrosine | Chemical | -0.26 (-0.41,-0.12) | 0.024 | 6 |
| sphingomyelin (d18:2/14:0, d18:1/14:1)* | Sphingomyelins | 0.26 (0.12,0.40) | 0.024 | 4 |
| bilirubin degradation product, C17H18N2O4 (2) | Hemoglobin and Porphyrin Metabolism | -0.26 (-0.40,-0.12) | 0.024 | 1 |
| bilirubin degradation product, C17H18N2O4 (3) | Hemoglobin and Porphyrin Metabolism | -0.26 (-0.40,-0.12) | 0.027 | 1 |
| bilirubin degradation product, C17H18N2O4 (1) | Hemoglobin and Porphyrin Metabolism | -0.26 (-0.40,-0.11) | 0.027 | 1 |
| 3-hydroxy-2-ethylpropionate | Leucine, Isoleucine and Valine Metabolism | -0.25 (-0.39,-0.11) | 0.031 | 7 |
| glycocholenate sulfate* | Secondary Bile Acid Metabolism | -0.25 (-0.39,-0.11) | 0.032 | 9 |
| cortisone | Corticosteroids | -0.25 (-0.39,-0.11) | 0.033 | 10 |
| 3-hydroxydecanoylcarnitine | Fatty Acid Metabolism (Acyl Carnitine, Hydroxy) | -0.24 (-0.38,-0.10) | 0.040 | 3 |
| succinimide | Chemical | -0.24 (-0.38,-0.10) | 0.040 | 1 |
| bilirubin degradation product, C17H20N2O5 (1) | Hemoglobin and Porphyrin Metabolism | -0.24 (-0.38,-0.10) | 0.044 | 1 |
| bilirubin degradation product, C17H20N2O5 (2) | Hemoglobin and Porphyrin Metabolism | -0.24 (-0.38,-0.10) | 0.047 | 1 |
| metabolonic lactone sulfate | Partially Characterized Molecules | 0.23 (0.09,0.38) | 0.049 | 12 |
| 3-hydroxyoctanoylcarnitine (1) | Hemoglobin and Porphyrin Metabolism | -0.23 (-0.37,-0.09) | 0.049 | 3 |
| pregnenolone sulfate | Pregnenolone steroids | -0.23 (-0.37,-0.09) | 0.049 | 14 |

Model fitted: metabolite ~ BMI.score.group (low BMI group as reference group). Model run on rank-based normal transformation (RNT) metabolite data. Beta represents change in normalised standard deviation units. Metabolites ordered by their Benjamini-Hochberg (BH) adjusted *p*-values from the lowest to the highest.
BH = Benjamini-Hochberg; CI = confidence interval.
\*: Indicates a compound that has not been confirmed based on a standard.
\*\*: Representative metabolite for clusters consisting of more than one metabolite.
<sup>a</sup>: Metabolite clusters assigned by iPVs (clusters 8, 11, 13 and 15 contain a single unidentified metabolite each)

Of the 68 xenobiotic metabolites (expressed as P/A traits) tested in Model 2, one metabolite, 2-acetamidophenol sulfate, had evidence for association with BMI score group. This metabolite was present less often in the plasma samples of individuals from the high BMI GS group (OR=0.59, 95%CI: 0.44,0.79, BH *p*=0.03). For full results of 68 metabolites from the logistic analysis see **Supplementary Table S4**.

### Sensitivity analyses

Characterisation of the score groups indicated some association with mother’s and mother’s partner’s social class. Therefore, in sensitivity analyses, these variables were fitted alongside score group in a multivariate model for the 29 metabolites with BH *p*<0.05 in the primary analysis. Score group effect estimates from the multivariate model (**Supplementary Table S5**) were similar to those from Model 1 (Pearson’s correlation, r=0.99).

Metabolite correlation analysis grouped the 29 metabolites associated with score group in Model 1 into 15 clusters, each with a representative metabolite (**Table 2**). The largest cluster consisted of 11 biochemicals including two forms of bilirubin, seven bilirubin degradation products, biliverdin and succinimide. There were 12 single metabolite clusters. Of the 15 representative metabolites, 12 were associated with measured BMI (p<0.05) in a multivariate linear model with BMI and BMI score group fitted alongside age and sex, whilst 14 had effect estimates that were directionally concordant with their BMI score group association as derived in Model 1 (**Supplementary Table S6** and **Supplementary Figure S4)**.

Lower plasma levels of bilirubin degradation product (C16H18N2O5 (1)) (beta=-0.02, 95%CI: -0.033,-0.006, *p*=5.94×10^−3^), hippurate (beta=-0.02, 95%CI: -0.033,-0.008, *p*=1.37×10^−3^), perfluorooctanesulfonate (PFOS) (beta=-0.02, 95%CI: -0.036,-0.012, *p*=1.03×10^−4^), tridecenedioate (C13:1-DC)* (beta=-0.05, 95%CI: -0.062,-0.035, *p*=7.54×10^−12^) and cortisone (beta=-0.06, 95%CI: -0.079,-0.050, *p*=3.28×10^−17^) as well as higher levels of sphingomyelin (d18:2/16:0, d18:1/16:1)* (beta=0.07, 95%CI: 0.049,0.080, *p*=3.10×10^−16^) and metabolonic lactone sulfate (beta=0.10, 95%CI: 0.077,0.120, *p*=4.74×10^−18^) were associated with higher measured BMI and showed the same direction of association with measured BMI in both recall groups (a direction that was also concordant with the BMI score group association from the main analysis) (**Figure 5 and Supplementary Table S6**). Fitting an interaction term (measured BMI * BMI score group) in the model provided some evidence to support a difference in the measured BMI effect by BMI score group for bilirubin degradation product, C16H18N2O5 (1) (*p*=0.048) and tridecenedioate (C13:1-DC)* (*p*=0.046). There was less evidence to support an association of measured BMI with levels of 3-hydroxy-2-ethylpropionate, pregnenolone sulfate, O-sulfo-L-tyrosine and glycocholenate sulfate (**Supplementary Figure S5 and Supplementary Table S6**). Plots for the representative metabolites not shown in **Figure 5** can be found in **Supplementary Figure S5**.

**Figure 5.**
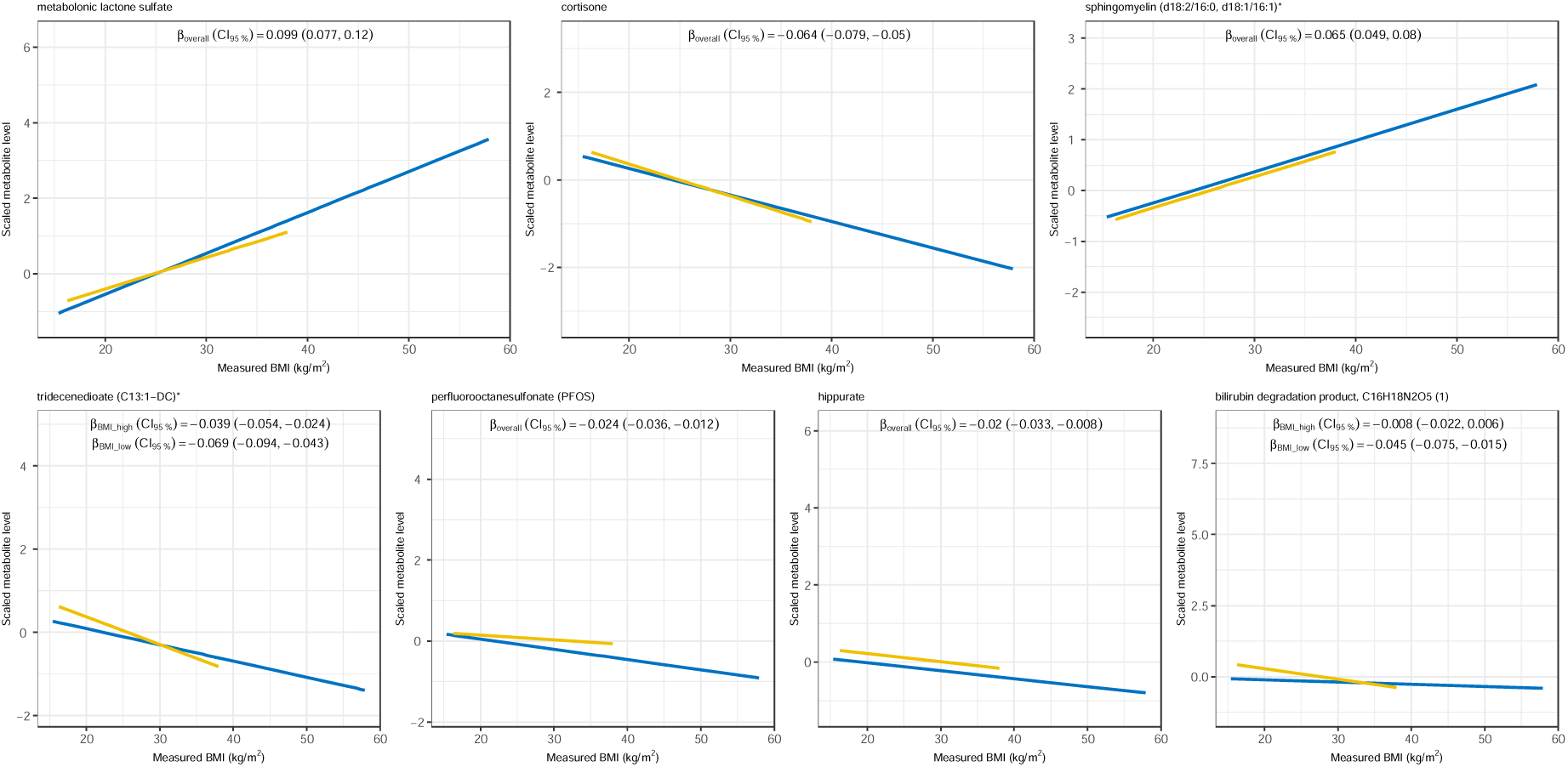
Relationship between selected BMI score group associated metabolites and measured BMI. Based on measured BMI at age 24 years clinic visit. Yellow = low BMI score group; blue = high BMI score group. *β*_*overall*_ is the measured BMI effect (CI_95%_ = 95% confidence interval), extracted from multivariate linear model fitted in all individuals [metabolite ∼ BMI + BMI.score.group + sex + age]. Where there was evidence that including an interaction term improved the fit of the model, the measured BMI effect (adjusted for age and sex) is given for each BMI score group separately (*β*_*BMI_high*_, *β*_*BMI_low*_). In the plots, solid lines denote the predicted univariate within score group relationship between BMI and metabolite with a 95% confidence interval denoted by shading.

## Discussion

In this study, we characterised the metabolic profile associated with low/high genetic liability for higher BMI using an RbG framework. The mean difference in BMI between the low and high BMI score groups increased from early childhood, reaching a maximum of 2.8 kg/m^2^ (95% CI: 2.1, 3.4) at time of sampling when individuals were on average 24.5 years of age. This observation reflects differences in the ability of the GS to capture variation in BMI at different ages as shown previously in the same cohort (19). We identified 29 metabolites associated with BMI GS group allocation. In most cases, associated metabolites were seen at lower levels in the high BMI score group, with the largest effects seen for bilirubin, hippurate and tridecenedioate. Two sphingomyelin metabolites were seen at increased abundance in the high BMI score group. The potential relevance of a selection of these metabolites to health and disease is explored in **Supplementary Table S7**.

Conventionally, MR and RbG approaches are an attempt at isolating the causal contribution of modifiable exposures, such as BMI, to chosen outcomes. However, when the outcome is also a biological intermediate that may itself be directly proxied by (elements of) the genetic predictor used to capture variance in the exposure, as is the case with metabolites, the assumptions underpinning the causal inference no longer hold. Holmes and Davey Smith have previously described seven scenarios which could underpin observed associations between a genetic risk score for disease and potential biomarkers (20). The scenarios, all of which we propose could equally apply to the present study, include both ‘real’ GRS-to-trait associations (which are therefore informative with respect to underlying biology) and associations that represent potential artifacts. Whilst others have concluded that the majority of metabolic perturbations seen in obesity are a response to increased adiposity itself rather than shared genetic mechanisms (21), our results and those of Hsu *et al*. (22), suggest forward causal connections (i.e., metabolites involved in pathways leading to changes in BMI) may exist. Whilst both sets of metabolic pathways, cause and effect, may be informative with respect to predicting the risk of developing obesity-associated comorbidities, only the former is likely to be of therapeutic relevance for the prevention of obesity. It is within this context that we go on to discuss our findings in more detail.

Bilirubin showed the strongest association with BMI score group allocation and, together with its associated degradation products, formed the largest cluster of associated metabolites. Bilirubin is a key component of the heme catabolic pathway and a potent antioxidant, and in this study was found in lower abundance in the high BMI score group. Although bilirubin is not amongst the metabolites most commonly associated with BMI and adiposity (23), circulating levels of the metabolite have previously been found to be associated both with adiposity and indicators of metabolic health (in a concordant direction to what we observe) in an observational setting (21, 24–27). Recent experimental work, including animal studies, support an active role for bilirubin in improving cardiorenal and metabolic dysfunction, pointing to a range of potential mechanisms including activation of nuclear receptors for burning fat (28) and the reduction of inflammation in adipose tissues (based on biliverdin administration to high-fat diet-induced obese mice) (29). However, results from MR studies aimed at improving causal inference in the role of bilirubin in a number of diseases have yet to provide robust evidence of a causal contribution (30–35).

Elevated levels of branched chain amino acids (BCAAs) (leucine, isoleucine and valine), as well as some of their tissue metabolites, have been consistently detected in individuals with obesity (23). There is evidence from MR that supports a causal effect of BMI on circulating BCAA (based on N=12,644) (26) as well as downstream effects of BCAA on disease such as type 2 diabetes (36). In this study, we saw little evidence for higher levels of these metabolites in the high BMI score group as compared to the low BMI group (Model 1 betas range from 0.015 to 0.073 with unadjusted *p*-values from 0.32 to 0.84). Whilst this may seem at odds with existing MR evidence, it is not totally unexpected given the level of inconsistency in the wider literature. For instance, studies conducted in children have failed to observe a positive relationship between BMI and BCAA (37, 38) whilst results from a bidirectional MR provided evidence for a causal effect of valine on BMI (22). However, the apparent instrument-dependent nature of the findings in the latter (22) point to heterogeneity in the underlying biology. Levels of circulating BCAA are also known to be influenced by dietary intake (39) and a link has been proposed between the obesity-related rise in circulating BCAA and a decline in their catabolism in adipose tissue (40, 41), with further evidence suggesting that this could be tissue specific (42). Moreover, 3-hydroxy-2-ethylproprionate (a metabolite annotated to the ‘Leucine, isoleucine and valine metabolism’ sub-pathway and which is a product of isoleucine catabolism (43)) was observed to associate with BMI GS group but not with measured BMI. One potential explanation for this given previously reported associations of 3-hydroxy-2-ethylproprionate with muscle cross-sectional area (i.e., with body composition) (44), is that the association seen with score group is underpinned by differences in lean mass between the groups. However, we are not well-powered to investigate this hypothesis within the current study.

We observed associations between BMI score group and the levels of potentially diet-related metabolites, including hippurate and PFOS. Hippurate, or hippuric acid, a glycine conjugate of benzoic acid, is synthesised in the liver and kidney (45). The benzoic acid component is derived mainly via microbial and mammalian co-metabolism of large polyphenolic molecules contained in, for example, fruits and vegetables, to a range of simpler aromatics that are then further metabolised to benzoic acids (45). We observed lower levels of hippurate in individuals in the high BMI GS group, concordant with a previous identified associations between hippurate levels and visceral body fat mass (27, 46). Previous literature combining data on diet intake, visceral fat mass and gut microbial profiling, suggests the association of circulating (and urine excreted) levels of hippurate with adiposity and related health outcomes (45), is likely to be the result of a complex interaction between diet intake, microbiome diversity and composition, and adipose tissue function (27, 47). Individuals in the high BMI score group also had lower plasma PFOS levels. PFOS, as an anthropogenic organic pollutant with chemical and thermal stability, has been detected in drinking water and the diet (especially in fish and crustaceans) in multiple countries and areas across the globe and has a global toxic effect on human health (see **Supplementary Table S7**). Furthermore, several of the xenobiotics which appear to be present at different rates in the two groups (albeit not meeting our stringent threshold for association) may also be biomarkers of food consumption (e.g., acesulfame, betonicine, theanine).

The associations observed between BMI score groups and these metabolites suggest that at least some of the genetic predisposition to increased BMI may be conveyed either via dietary choices or through differences in nutrient metabolism. Others have taken the fact that many genes associated with high BMI appear to be highly expressed in the central nervous system (48) (with some also having been linked to appetite regulation), as evidence that genetic susceptibility to obesity is partly attributable to appetitive phenotypes (49). Studies of the association between BMI-associated GS and appetite and satiety traits (50, 51) along with results of the current study provide some support for this hypothesis. However, behavioural traits such as these are known to be particularly at risk from bias even in an MR (and likely RbG) setting (52), where population stratification (53) or complex genetic effects not accounted for in these study designs, for example, dynastic effects (54) can be problematic given the underlying assumptions of these methods.

The potential for reintroduction of such confounding effects (and pleiotropy) appears to increase with the number of variants included in the GS, either through the lowering of the p-value threshold used for selecting SNPs or through increasing the power of association analyses such that ever smaller effects are detected with sufficient statistical certainty (15). In this study, the weak correlation between score group and social class suggests some residual confounding (due to population stratification) may be present. However, given the consistency in the effect estimates after adjusting for maternal and paternal social class, we believe the effects of any such confounding on our results to be small. In this study, where metabolites showed an association with BMI score group, we also estimated their association with measured BMI. Whilst in general we saw good concordance of effects estimated in the two analyses, given our sampling frame, the observational estimates cannot necessarily be extrapolated to the wider (unselected) population. Where we see inconsistency, this could be the result of different key sources of bias effecting the different analytical strategies.

In conclusion, we used an innovative RbG study design to identify metabolites whose relative abundance varies with a genetic predisposition to increased BMI. We hypothesise that these differences may reflect gene-derived perturbations to biological pathways relevant to weight gain or be consequences of higher BMI itself. To provide a definitive answer to the question of the role of metabolites (and related biological pathways) in obesity, results from several different approaches with unrelated sources of bias, including challenge/intervention studies, need to be integrated. In doing so, we can begin to understand the role of different metabolic pathways in weight gain and related morbidity and partition metabolites according to whether they are likely to be causative in nature and/or reflective of metabolic changes that occur after increased adiposity.

## Acknowledgements

Avon Longitudinal Study of Parents and Children (ALSPAC): We are extremely grateful to all the families who took part in this study, the midwives for their help in recruiting them, and the whole ALSPAC team, which includes interviewers, computer and laboratory technicians, clerical workers, research scientists, volunteers, managers, receptionists and nurses. Individual participant data (phenotype and metabolite) that underline the results reported in this article will be available to researchers upon approval by the ALSPAC Executive Committee.

