## Supporting Information for "A multi-variant recall-by-genotype study of the metabolomic signature of body mass index"

**Fang *et al*.**

### Supplementary Methods

#### Avon Longitudinal Study of Parents and Children (ALSPAC): Cohort summary

##### Description of study numbers

Pregnant women resident in Avon, UK with expected dates of delivery 1st April 1991 to 31st December 1992 were invited to take part in the study. The initial number of pregnancies enrolled is 14,541 (for these at least one questionnaire has been returned or a “Children in Focus” clinic had been attended by 19/07/99). Of these initial pregnancies, there was a total of 14,676 foetuses, resulting in 14,062 live births and 13,988 children who were alive at 1 year of age.

When the oldest children were approximately 7 years of age, an attempt was made to bolster the initial sample with eligible cases who had failed to join the study originally. As a result, when considering variables collected from the age of seven onwards (and potentially abstracted from obstetric notes) there are data available for more than the 14,541 pregnancies mentioned above. The number of **new pregnancies** not in the initial sample (known as Phase I enrolment) that are currently represented on the built files and reflecting enrolment status at the age of 24 is 913 (456, 262 and 195 recruited during Phases II, III and IV respectively), resulting in an additional 913 children being enrolled. The phases of enrolment are described in more detail in the cohort profile paper and its update (see footnote 4 below). The total sample size for analyses using any data collected after the age of seven is therefore 15,454 pregnancies, resulting in 15,589 foetuses. Of these 14,901 were **alive at 1 year of age**.

A 10% sample of the ALSPAC cohort, known as the **Children in Focus (CiF) group**, attended clinics at the University of Bristol at various time intervals between 4 to 61 months of age. The CiF group were chosen at random from the last 6 months of ALSPAC births (1432 families attended at least one clinic). Excluded were those mothers who had moved out of the area or were lost to follow-up, and those partaking in another study of infant development in Avon.

##### Data collection

Study data were collected and managed using REDCap electronic data capture tools hosted at the University of Bristol (1, 2). REDCap (Research Electronic Data Capture) is a secure, web-based software platform designed to support data capture for research studies, providing 1) an intuitive interface for validated data capture; 2) audit trails for tracking data manipulation and export procedures; 3) automated export procedures for seamless data downloads to common statistical packages; and 4) procedures for data integration and interoperability with external sources.

Please note that the study website contains details of all the data that is available through a fully searchable data dictionary and variable search tool (3).

##### Ethical approvals

Ethical approval for the study was obtained from the ALSPAC Ethics and Law Committee and the Local Research Ethics Committees (4). Consent for biological samples has been collected in accordance with the Human Tissue Act (2004). Informed consent for the use of data collected via questionnaires and clinics was obtained from participants following the recommendations of the ALSPAC Ethics and Law Committee at the time.

##### Genotyping

ALSPAC children were genotyped using the Illumina HumanHap550 quad chip genotyping platforms by 23andme subcontracting the Wellcome Trust Sanger Institute, Cambridge, UK and the Laboratory Corporation of America, Burlington, NC, US. The resulting raw genome-wide data were subjected to standard quality control methods. Individuals were excluded on the basis of gender mismatches; minimal or excessive heterozygosity; disproportionate levels of individual missingness (>3%) and insufficient sample replication (IBD < 0.8). Population stratification was assessed by multidimensional scaling analysis and compared with Hapmap II (release 22) European descent (CEU), Han Chinese, Japanese and Yoruba reference populations; all individuals with non-European ancestry were removed. SNPs with a minor allele frequency of < 1%, a call rate of < 95% or evidence for violations of Hardy-Weinberg equilibrium (P < 5E-7) were removed. Cryptic relatedness was measured as proportion of identity by descent (IBD > 0.1). Related subjects that passed all other quality control thresholds were retained during subsequent phasing and imputation. 9,115 subjects and 500,527 SNPs passed these quality control filters.

ALSPAC mothers were genotyped using the Illumina human660W-quad array at Centre National de Génotypage (CNG) and genotypes were called with Illumina GenomeStudio. PLINK (v1.07) was used to carry out quality control measures on an initial set of 10,015 subjects and 557,124 directly genotyped SNPs. SNPs were removed if they displayed more than 5% missingness or a Hardy-Weinberg equilibrium P value of less than 1.0e-06. Additionally, SNPs with a minor allele frequency of less than 1% were removed. Samples were excluded if they displayed more than 5% missingness, had indeterminate X chromosome heterozygosity or extreme autosomal heterozygosity. Samples showing evidence of population stratification were identified by multidimensional scaling of genome-wide identity by state pairwise distances using the four HapMap populations as a reference, and then excluded. Cryptic relatedness was assessed using a IBD estimate of more than 0.125 which is expected to correspond to roughly 12.5% alleles shared IBD or a relatedness at the first cousin level. Related subjects that passed all other quality control thresholds were retained during subsequent phasing and imputation. 9,048 subjects and 526,688 SNPs passed these quality control filters.

We combined 477,482 SNP genotypes in common between the sample of mothers and sample of children. We removed SNPs with genotype missingness above 1% due to poor quality (11,396 SNPs removed) and removed a further 321 subjects due to potential ID mismatches. This resulted in a dataset of 17,842 subjects containing 6,305 duos and 465,740 SNPs (112 were removed during liftover and 234 were out of HWE after combination). We estimated haplotypes using ShapeIT (v2.r644) which utilises relatedness during phasing. We obtained a phased version of the 1000 genomes reference panel (Phase 1, Version 3) from the Impute2 reference data repository (phased using ShapeIt v2.r644, haplotype release date Dec 2013). Imputation of the target data was performed using Impute V2.2.2 against the reference panel (all polymorphic SNPs excluding singletons), using all 2186 reference haplotypes (including non-Europeans).

This gave 8,237 eligible children and 8,196 eligible mothers with available genotype data after exclusion of related subjects using cryptic relatedness measures described previously.

#### Genetic score derivation

Samples were selected for inclusion in the study based on a genetic score (GS) for body mass index (BMI). We used publicly available summary statistics from the combined GIANT and UK Biobank meta-GWAS analysis of BMI (5) downloaded from: <https://portals.broadinstitute.org/collaboration/giant/index.php/GIANT_consortium_data_files#BMI_and_Height_GIANT_and_UK_BioBank_Meta-analysis_Summary_Statistics> (last accessed: October 2018), to calculate a weighted GS. Specifically, we included 940 of the 941 SNPs listed in the paper as near-independent SNPs associated with BMI (at a revised genome-wide significance threshold of P<1×10^−8^) following an approximate conditional and joint multiple-SNP (COJO) analysis that considers LD between SNPs at a given locus (6). Within the set of 941 SNPs are 656 main associations and 285 secondary associations. The variant rs1000096 was the only SNP from the 941 not included in our GS (due to a coding error).

Genetic scores were calculated for all ALSPAC G1 with genetic data (N=8,953). To generate the GS, first, genotypes for the subset of 940 SNPs required were extracted from the full (imputed) ALSPAC dataset using QCTOOL v2. The .gen files produced for each chromosome were then concatenated to make a single .gen file. The genotype data was then converted to dosage format, again, using QCTOOL v2 and manually reformatted to match the dosage format required for PLINK (including a swap of the reference allele). Finally, the PLINK --score function was used with the options: ‘sum’, ‘double-dosage’ and ‘include-cnt’ (7).

#### Sample selection

Individuals were ordered by their GS and those within the top and bottom 30% of the distribution crossmatched against a list of individuals who both attended the age 24 years clinic visit and had a stored plasma sample available for analysis. This left N=738 individuals from the top 30% and N=806 from the bottom 30% in the eligible set. A scheme was then devised to select those with the most extreme GS from the two groups (highest and lowest) whilst maintaining some degree of balance (in extremity) across the groups. First, individuals in the top set were assigned a rank, starting with the individual with the highest (most extreme) GS. Conversely, individuals in the bottom set were assigned a rank starting with the individual with the lowest GS. Based on these ranks and starting from rank 1, matched (on rank) pairs of individuals were selected for inclusion until 240 pairs had been selected. A further 138 individuals per set were then selected for inclusion, starting with those with the highest rank (i.e., the most extreme) in each set who had not yet been selected as part of a pair. This scheme resulted in an approximately balanced (by rank) sample of N=378 with the highest GS in the top set and N=378 with the lowest GS in the bottom set being selected for inclusion in the study. In addition, four duplicate samples were included (two per group). In total, 760 samples were sent to Metabolon for metabolomics analysis.

#### Sample collection

Blood samples were collected at the age 24 years clinic visit from all ALSPAC G1 individuals who provided informed consent. Participants were instructed to fast for a minimum of eight hours prior to their appointment and 90% complied (the remainder having consumed food within the preceding eight hours). During clinic appointments, blood was drawn into 10ml K2E (K2EDTA) tubes (VACUETTE®) that were centrifuged at 3500rpm for 10 minutes at 4–5°C. Plasma was then transferred to 200ul aliquots and stored at −80°C. Samples were processed and frozen within 90 minutes of collection, where possible. Selected samples were shipped on dry ice to Metabolon, Inc. for untargeted metabolomics analysis using established protocols.

#### Derivation of metabolite data by Metabolon, Inc.

The methodological details provided herein are as supplied by Metabolon, Inc.

##### Sample preparation

Samples were prepared using the automated MicroLab STAR® system from Hamilton Company. Several recovery standards were added prior to the first step in the extraction process for QC purposes. Proteins were precipitated with methanol under vigorous shaking for 2 min (Glen Mills GenoGrinder 2000) followed by centrifugation. The resulting extract was divided into five fractions: two for analysis by two separate reverse phase (RP)/UPLC-MS/MS methods with positive ion mode electrospray ionization (ESI), one for analysis by RP/UPLC-MS/MS with negative ion mode ESI, one for analysis by HILIC/UPLC-MS/MS with negative ion mode ESI, and one for backup. Samples were placed briefly on a TurboVap® (Zymark) to remove the organic solvent. The sample extracts were stored overnight under nitrogen before preparation for analysis.

##### Quality assurance / Quality control

Three types of controls were used when analyzing the experimental samples: a pooled matrix sample generated by taking a small volume of each experimental sample (or alternatively, use of a pool of well-characterized human plasma); extracted water samples; and a cocktail of QC standards. Instrument variability was determined by calculating the median relative standard deviation (RSD) for the standards that were added to each sample prior to injection into the mass spectrometers. Overall process variability was determined by calculating the median RSD for all endogenous metabolites (i.e., non-instrument standards) present in 100% of the pooled matrix samples. Experimental samples were randomized across the platform run with QC samples spaced evenly among the injections.

##### Ultrahigh Performance Liquid Chromatography-Tandem Mass Spectroscopy (UPLC-MS/MS)

All methods utilized a Waters ACQUITY ultra-performance liquid chromatography (UPLC) and a Thermo Scientific Q-Exactive high resolution/accurate mass spectrometer interfaced with a heated electrospray ionization (HESI-II) source and Orbitrap mass analyzer operated at 35,000 mass resolution. The sample extract was dried then reconstituted in solvents compatible to each of the four methods. Each reconstitution solvent contained a series of standards at fixed concentrations to ensure injection and chromatographic consistency. One aliquot was analyzed using acidic positive ion conditions, chromatographically optimized for more hydrophilic compounds. In this method, the extract was gradient eluted from a C18 column (Waters UPLC BEH C18-2.1x100 mm, 1.7 µm) using water and methanol, containing 0.05% perfluoropentanoic acid (PFPA) and 0.1% formic acid (FA). Another aliquot was also analyzed using acidic positive ion conditions, however it was chromatographically optimized for more hydrophobic compounds. In this method, the extract was gradient eluted from the same afore mentioned C18 column using methanol, acetonitrile, water, 0.05% PFPA and 0.01% FA and was operated at an overall higher organic content. Another aliquot was analyzed using basic negative ion optimized conditions using a separate dedicated C18 column. The basic extracts were gradient eluted from the column using methanol and water, however with 6.5mM Ammonium Bicarbonate at pH 8. The fourth aliquot was analyzed via negative ionization following elution from a HILIC column (Waters UPLC BEH Amide 2.1x150 mm, 1.7 µm) using a gradient consisting of water and acetonitrile with 10mM Ammonium Formate, pH 10.8. The MS analysis alternated between MS and data-dependent MSn scans using dynamic exclusion. The scan range varied slighted between methods but covered 70-1000 m/z.

##### Data extraction and compound identification

Raw data was extracted, peak-identified and QC processed using Metabolon’s hardware and software. Compounds were identified by comparison to library entries of purified standards or recurrent unknown entities. More than 3300 commercially available purified standard compounds have been acquired and registered into LIMS for analysis on all platforms for determination of their analytical characteristics. Additional mass spectral entries have been created for structurally unnamed biochemicals, which have been identified by virtue of their recurrent nature (both chromatographic and mass spectral).

##### Metabolite Quantification and Data Normalization

Peaks were quantified using area-under-the-curve. A data normalization step was performed to correct variation resulting from instrument inter-day tuning differences.

#### Definitions of variables

##### Adiposity traits

**Weight (kg) and BMI (kg/m^2^):** Weight (kg) and height (m) measures were assessed at all clinic visits. Standing height was measured to the nearest millimetre using a wall-mounted stadiometer. Weight was measured to the nearest 0.1kg using Tanita TBF-401A electronic body composition scales (or electronic bathroom scales, if the participant had a pacemaker). Body mass index (BMI) was calculated as [weight (kg)] / [height (m)^2^]. In addition to the height and weight measures obtained at ALSPAC clinics, weight and BMI measures derived from other data sources (specifically mother-completed questionnaires and health visitor records) between the ages of four months and ten years were included in analyses (8).

**Body composition phenotypes:** Dual emission x-ray absorptiometry (DXA) was used to measure fat mass, muscle mass and bone density during the age 24 years clinic visit. Total body lean mass (g) and total body fat mass (g) were determined by full body DXA scans and transformed to kg prior to analysis.

##### Potential confounders

Data were extracted for several phenotypic correlates of observed BMI to check for associations with score group and evaluate the potential for them to act as confounders in the primary analysis. Several phenotypes were selected measured in the G1 themselves (sex, age at the sample collection clinic (in weeks) and moderate to vigorous physical activity (MVPA) (minutes per day)), their mothers (parity, highest education, alcohol drinking status, smoking history, and social class) or their mother’s partners (social class). Fasting status (binary) and time since last food/drink (in hours) were also evaluated as potential technical confounders.

**Age (in weeks):** Participant’s age was recorded as their age at clinic visit. At different clinic visits, age was recorded in different units, including weeks, months and years. If age in weeks data were available, the age was used in the present study directly. Otherwise, age data were converted to weeks by multiplying age in months data by 4.34524.

**Moderate to Vigorous Physical Activity (MVPA) (minutes per day):** At the age 24 years clinic, participants were asked to wear an ActiGraph GT3X+ accelerometer device after the clinic visit for four consecutive days which were part of a “normal week” instead of doing anything unusual. An objective measurement of human activity was measured by the device. MVPA referred to activities with a counts per minute (cpm) of more than 2020. MVPA minutes used in the present study was defined as the average of MVPA minutes per day over all days the device was worn.

**Parental phenotypes:** Phenotypic data relating to the participant’s mother (parity, smoking history, alcohol drinking status, highest education, and social class), and her partner’s social class were collected via questionnaires completed by the mother during pregnancy. Parity is defined as the number of previous pregnancies resulting in either a livebirth or a stillbirth. Mother’s smoking history is defined as the mother’s answer to “Have you ever been a smoker?”, being either “Yes” or “No”. Mother’s alcohol drinking status records the alcohol consumption frequency of the mother before the current pregnancy with six categories (“never”, “<1 glass per week”, “1+ glasses per week”, “1-2 glasses per day”, “3-9 glasses per day”, “10+ glasses per day”). Mother’s highest education qualification is a categorical variable with five values (“CSE/none”, “Vocational”, “O level”, “A level” and “Degree”). Mother’s and partner’s social classes are derived variables generated based on the type of industry and level of their occupation with seven different values.

**Fasting status:** Two variables, recorded during the age 24 years clinic visit, were used to assess fasting status. Participants were instructed to fast for a minimum of eight hours before attending the clinic during which blood sampling was undertaken. Therefore, during the clinic visit, participants were asked when they last ate or drank and two variables derived: (1) a continuous phenotype – time since last food/drink (in hours); a binary phenotype indicating whether food/drink had been consumed in the last 8 hours or not, i.e., fasting status.

#### Metabolite data pre-analysis processing

##### Metabolite data processing pipeline

We processed the raw (original scale) data received from Metabolon (N=760 samples) in preparation for statistical analysis using an in-house pipeline developed in R (9) (a pre-release version of the R package ‘metaboprep’ currently available from: <https://github.com/MRCIEU/metaboprep>). First, all metabolites designated (by Metabolon) as xenobiotics were temporarily removed from the dataset. The reason for this is that xenobiotics are metabolites not produced by the body, such as drug compounds, and therefore can have very high rates of missingness, while still being critically informative to a study. Next, we screened for particularly poorly performing samples and metabolites, defined as those with >80% missing data. Missingness was then re-assessed based on remaining samples and metabolites and a more stringent missingness criteria of >20% applied (i.e., samples and metabolites with >20% missing values removed). No samples were excluded based on missingness at either threshold, whilst 211 metabolites were removed (22 with >80% missing and a further 189 with >20% missing). Sample quality was further assessed based on total peak area (TPA), calculated as the sum of all metabolite values measured in a sample, with a single sample whose TPA fell more than five standard deviations (SDs) from the mean being excluded. Finally, a principal component analysis (PCA) was conducted on the samples using a subset of approximately independent metabolites (see below for details) to identify potential outliers. Two samples positioned more than five SDs from the mean of the first and/or second principal components were considered outliers and excluded from subsequent analysis. Following these procedures, data for the xenobiotic metabolites were added back into the dataset. Further sample exclusions were applied based on consent withdrawals (N=3) and duplicate samples (N=4). In the case of the latter, samples with the least missing data from each pair were retained for analysis. After processing, 750 samples (377 from high score group and 373 from low score group) and 1005 metabolites remained.

Of the 1005 metabolites remaining after processing, 905 were (non-xenobiotic) metabolites with <20% missing data. In this dataset, missing data was imputed using a random-forest based method implemented in the missForest R package (10). Then, data for this set of metabolites were transformed by rank-based normal transformation (RNT) to ensure normality prior to statistical analysis. RNT was achieved using a modified version of the rntransform() function (from the GenABEL package) such that any tied values were randomly split (code is available from the moosefun git repository at: <https://github.com/hughesevoanth/moosefun>). Meanwhile, data for the 100 xenobiotic metabolites with >20% missing data were transformed to presence/absence (P/A) data, where 1 represents presence in a sample and 0 represents absence. Finally, xenobiotics present in less than 11 samples (N=32) were excluded from downstream analyses on the basis that any statistical analyses would not be robust.

##### Selection of features for principal component analysis

Prior to conducting the PCA to identify sample outliers, a subset of approximately independent features was selected as follows. Data were restricted to common metabolites such that only those that were (a) variable and (b) had at least 50 observations were included. A dendrogram was then constructed (stats package hclust() function, with method ‘complete’) based on a Spearman’s rho distance matrix (1-|Spearman’s rho|). A set of ‘k’ clusters (groups of similar metabolites) were identified based on a user-defined tree cut height 0.2 (equivalent to a Spearman’s rho of 0.8), using the function cutree() from the stats package. For each ‘k’ cluster the metabolite with the least missingness was then tagged as the representative metabolite for that cluster. Data for the subset of representative metabolites then underwent imputation with missing values imputed to the median and finally data standardised (z-transformed) such that the mean equals zero and the standard deviation equals one for each metabolite. These data were then used as input to the PCA analysis conducted as part of the data processing pipeline described above.

#### Extended methods relating to sensitivity analyses

##### Extension of primary association analyses

Model 1 was extended to a multivariate model in which any potential confounder that had previously been shown to be associated with score group was fitted as an independent fixed effect alongside score group. In addition to extracting the model coefficients, the variance explained by each of the fixed effects in the model (designated ‘VE’ in results files) was estimated by Type II ANOVA (‘Anova()’ function from the ‘car’ package in R (11)).

##### Metabolite correlation analysis

We took the subset of associated metabolites output from Model 1 and used a hierarchical clustering approach to identify redundancy in the data (i.e., where associated metabolites were highly correlated and likely representing the same biological signal). This was done based on the original (unimputed abundance) data and using the iPVs R package (12) with a tree cut height of 0.75 (where the value is equal to a dissimilarity of 1 - Spearman's rho). As well as assigning metabolites to clusters based on their similarity, this package uses principal variable analysis (PVA) to select the best representative metabolite of each cluster. The set of reduced ‘representative’ metabolites was used as the focus for the next steps.

##### Association of metabolites with measured BMI

A series of linear regression analyses were conducted to evaluate the direct association between measured BMI (at the age 24 years clinic) and the subset of BMI score group associated metabolites. Having established that the (associated) metabolite distributions were approximately normal, original (unimputed abundance) data was mean centred and scaled by the standard deviation (z-scored) and the overall association between measured BMI and metabolite level assessed with BMI score group, sex and age fitted as covariates in a multivariate linear model [metabolite ~ BMI + BMI.score.group + sex + age]. In order to investigate the consistency of the BMI effect across the two groups, the same model was also fitted with an interaction term [metabolite ~ BMI * BMI.score.group + sex + age] and within each BMI score group separately [metabolite ~ BMI + sex + age]. Robust standard errors generated with coeftest() from lmtest R package were reported because of heteroscedasticity. ANOVA tests were carried out to ascertain whether including the interaction effect improved the model fit. Variance explained by each fixed effect in the model was calculated as described previously.

### Supplementary Tables

#### Table S1. Between-group difference in BMI and weight from 4months to 24 years of age (in Excel file)

See separate Excel file.

#### Table S2. Association of potential confounders of primary analysis with BMI score group

| **Phenotype category** | **Statistical test** | **Phenotype** | **Mean difference (95% CI)^a^** | **p-value** |
| --- | --- | --- | --- | --- |
| Continuous | Student’s two-sample two-sided t-test | MVPA (minutes) | -1.02 (-10.1,8.02) | 0.82 |
|  |  | Age at clinic (weeks) | -0.05 (-5.85,5.76) | 0.99 |
|  |  | Time since last food at sampling (hours) | -0.21 (-0.70,0.28) | 0.40 |
| Categorical | Fisher’s exact test for count data | Mother’s highest education | na | 0.15 |
|  |  | Mother’s social class | na | 0.08 |
|  |  | Mother’s partner’s social class | na | 0.02 |
|  |  | Mother’s parity | na | 0.83 |
|  |  | Mother’s alcohol drinking status (frequency) | na | 0.29 |
| Binary |  | Mother’s smoking status | 0.87 (0.63,1.19) | 0.40 |
|  |  | Fasting status at sampling | 0.69 (0.40, 1.20) | 0.19 |

CI = confidence interval;

^a^ expressed as the mean difference in the high BMI score group as compared to the low BMI score group as the reference.

#### Table S3. Results of linear regression for 905 metabolites (in Excel file)

See separate Excel file.

#### Table S4. Results of logistic regression for 68 metabolites (in Excel file)

See separate Excel file.

#### Table S5. Results of linear regression adjusting for maternal and paternal social class for 25 metabolites associated with BMI score group in the primary analysis (in Excel file)

See separate Excel file.

#### Table S6. Results of linear regression of metabolite levels on observed BMI for 25 associated representative metabolites (in Excel file)

See separate Excel file.

#### Table S7. Literature summary by metabolite

| Metabolite name | Summary of findings | General information | Relevance to disease |
| --- | --- | --- | --- |
| Bilirubin | A total of 10 bilirubin related metabolites (including biliverdin) were found to be lower in individuals with high BMI genetic score. | Bilirubin is a key component of the heme catabolic pathway, in which the heme present in hemoproteins from red blood cell catabolism are oxygenated (by heme oxygenase) to biliverdin, and then reduced by biliverdin reductase to bilirubin. | bilirubin has been found to have a protective role against inflammation, hepatic diseases, diabetes, metabolic syndrome and obesity (13–16). |
| Sphingomyelin | Sphingomyelin (d18:2/16:0, d18:1/16:1) and sphingomyelin (d18:2/14:0, d18:1/14:1) showed higher levels in individuals from high BMI genetic score group. | Sphingomyelins are the most abundant class of sphingolipids in lipoproteins with essential roles in both maintaining plasma membrane structures and cellular signalling (17). Sphingomyelin consist of multiple species, with sphingosine (d18:1) being the most prominent long-chain base and palmitic acid (16:0) as the most common fatty acid component (17, 18). | It has been long recognised that high levels of plasma sphingomyelin promote atherogenesis, and it is an independent risk factor for coronary artery disease (19, 20). |

| Metabolite name | Summary of findings | General information | Relevance to disease |
| --- | --- | --- | --- |
| Hippurate | We observed lower levels of hippurate in individuals in the high BMI GS group and a corresponding negative association with measured BMI. | Hippurate, or hippuric acid, is a mammalian-microbial cometabolite synthesised by glycine conjugation of benzoic acid in the liver and kidney and from microbial metabolism of diet-derived polyphenols (21). | Higher levels of hippurate have previously been associated with higher fruit and whole grain intake, lower visceral fat mass and reduced risk of having metabolic syndrome features (22–24). |
| PFOS | We found individuals in the high BMI score group had lower plasma PFOS levels and observed a corresponding negative association with measured BMI. | PFOS is one of the many anthropogenic organic pollutant perfluoroalkyl and polyfluoroalkyl substances (PFASs) with chemical and thermal stability. They have been widely applied as industrial and commercial polymers and surfactants since 1950 (25). Despite most uses of PFOS having been phased out, banned or restricted under a number of UK, EU and international regulations, PFOS remains a widespread environmental contaminant (26). Studies published as recently as 2019 have confirmed the presence of PFOS in drinking water in multiple countries and areas across the globe (27) and the diet in Canada (28), Spain (29), the Netherlands (30) and Greece (31), especially in fish and crustaceans. | In humans, PFOS accumulates in multiple tissues and body fluids with enrichments in blood (32) and the liver (33). Molecular and animal studies have shown that PFOS has a global toxic effect on human health, including detrimental impacts on the liver, neuron system, reproductive system and immune system (34). |

| Metabolite name | Summary of findings | General information | Relevance to disease |
| --- | --- | --- | --- |
| Cortisone | We observed lower levels of cortisone among individuals in the high BMI score group and this association was supported in the analysis of measured BMI. | Cortisone is an inert metabolite of cortisol, a steroid hormone released by the adrenal cortex. Cortisone regenerates cortisol, and its levels are regulated by reversible enzyme shuttle between two subtypes of 11β-Hydroxysteroid dehydrogenase (11β-HSD) (35). | There is a long history of research into cortisol, the precursor of cortisone, and obesity (36). Whilst a cross-sectional study and review of the literature published in 2013 concluded there was no strong relationship between systemic cortisol and obesity (37), more recent work has pointed to associations between both cortisone and cortisol and specific fat depots (24). |
| O-sulfo-L-tyrosine | The level of O-sulfo-L-tyrosine was found to be lower in individuals with higher BMI genetic score, but it did not show strong association with measured BMI in both recall groups. | O-sulfo-L-tyrosine is a metabolite from the class of phenylalanine and derivatives. | Previous studies have identified an association between higher O-sulfo-L-tyrosine levels and reduced kidney function in chronic kidney disease (38, 39). |
| 3-hydroxy-2-ethylpropionate | Low levels of 3-hydroxy-2-ethylpropionate levels were observed in the high BMI score group, but it did not show strong association with measured BMI in both recall groups. | 3-hydroxy-2-ethylpropionate, or 2-ethylhydracrylate, is a metabolite involved in the metabolism of leucine, isoleucine and valine, the branched chain amino acids. | 3-hydroxy-2-ethylpropionate has been found to be positively associated with muscle cross-sectional areas and reversely associated with the proinflammatory cytokines tumour necrosis factor-α (TNF-α) (40). |
| Metabolite name | Summary of findings | General information | Relevance to disease |
| Glycocholenate sulfate* | Low levels of glycocholenate sulfate levels were observed in the high BMI score group, but it did not show strong association with measured BMI in both recall groups. | Glycocholenate sulfate is a circulating secondary bile acid. | Circulating glycocholenate sulfate level was found to be positively associated with incidence of atrial fibrillation (AF) risk in multiple studies (41, 42). Another study suggested it was inversely associated with high-grade glioma (43). |

PFOS - Perfluorooctane sulfonate;

*: Indicates a compound that has not been confirmed based on a standard.

### Supplementary Figures

#### Supplementary Figure S1. Mean differences in weight between the high and low BMI score groups.

Error bars represent the standard errors of mean differences in weight. Sample size ranges from 111 (at age 31 months) to 744 (at age 24 years). Test results are given for a Students (two-sample two-sided) t-test. ***: p-value < 0.001; **: p-value < 0.01; *: p-value < 0.

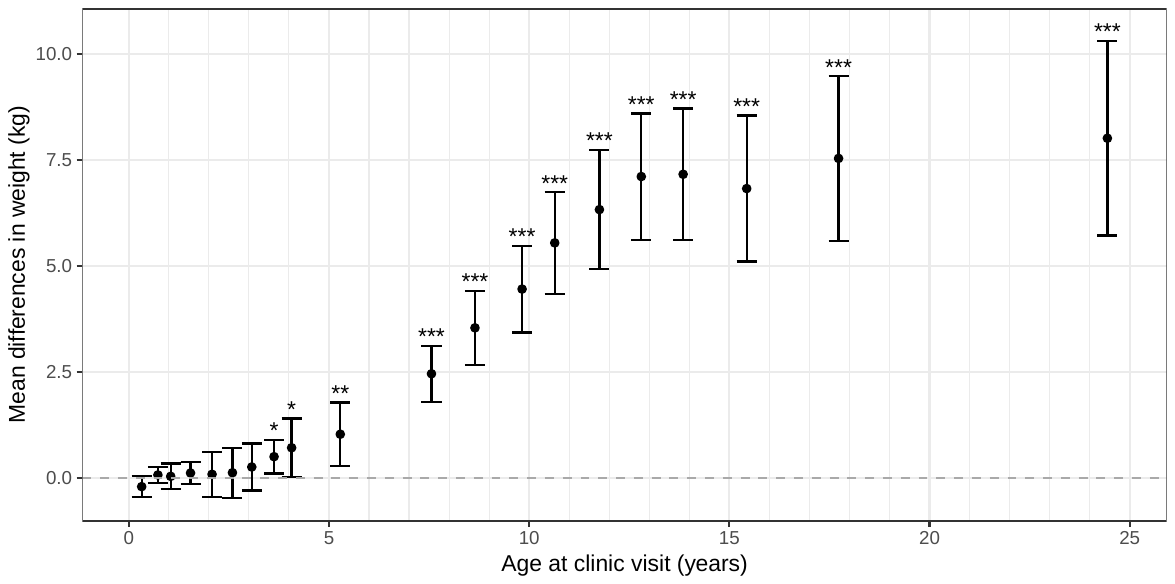

#### Supplementary Figure S2. Distribution across social class categories by BMI score group

| 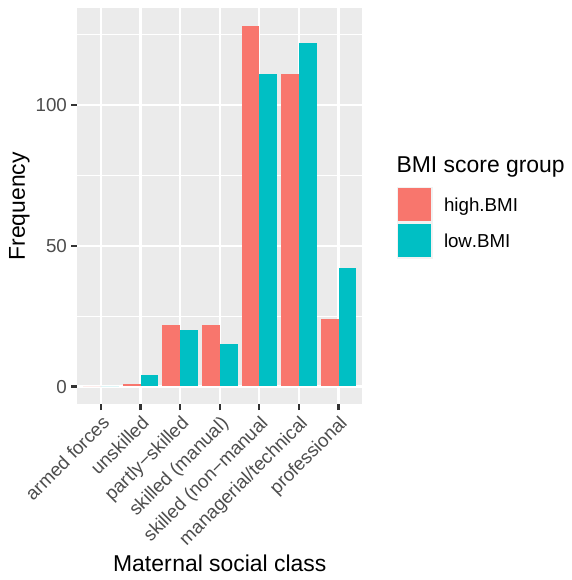 | 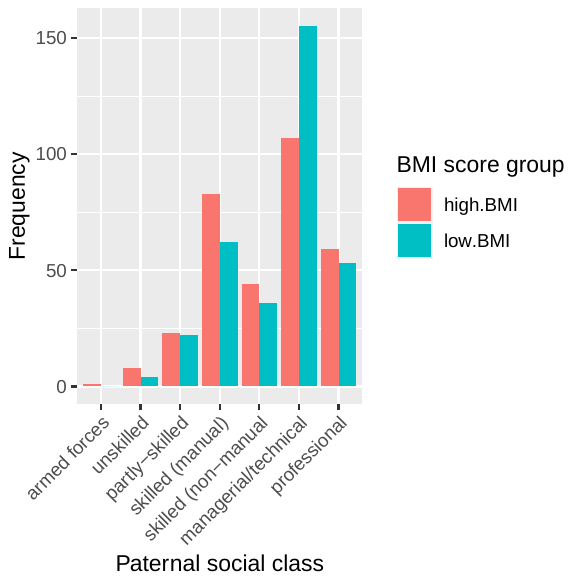 |
| --- | --- |

#### Supplementary Figure S3. Distribution of z-scored levels of BMI score group associated metabolites by group.

See separate PDF.

#### Supplementary Figure S4. Comparison of BMI score group effects estimated from Model 1 to BMI effect estimates based on measured BMI

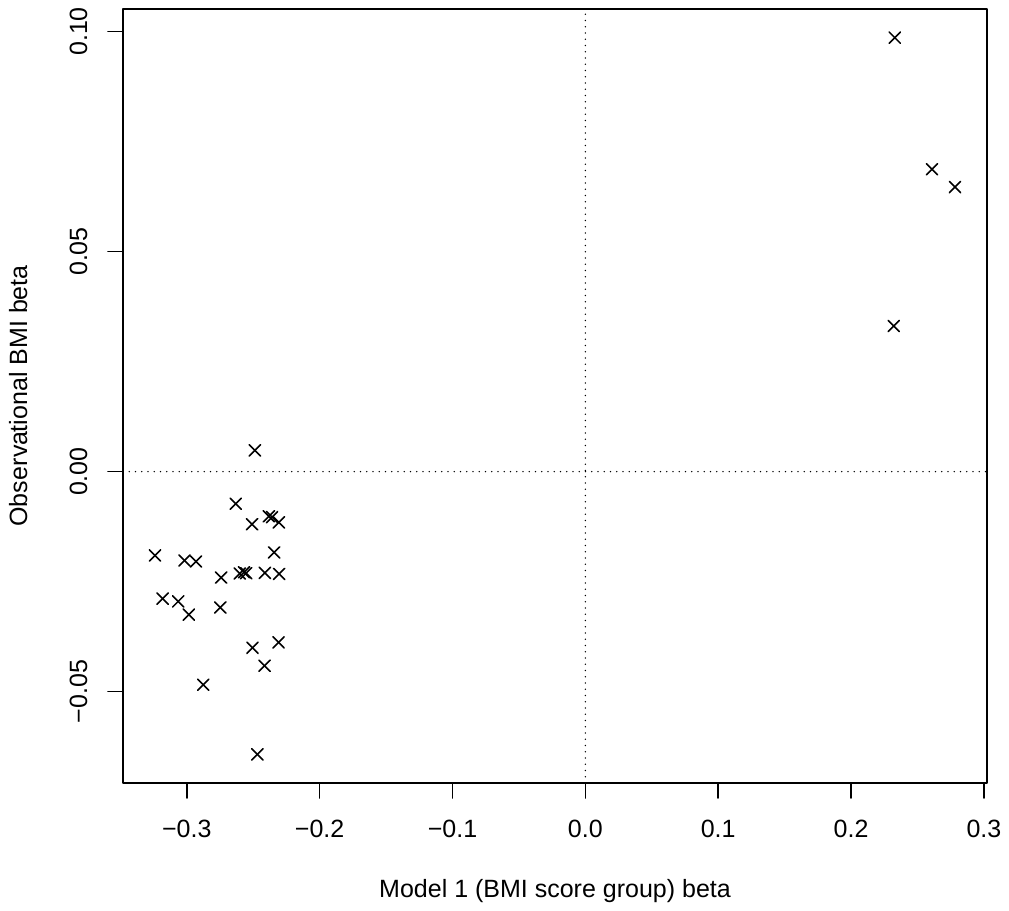

#### Supplementary Figure S5. Relationship between selected BMI score group associated metabolites and measured BMI.

Based on measured BMI at the age 24 years clinic. Yellow = low BMI score group; blue = high BMI score group. $\beta_{overall}$ is the measured BMI effect (CI_95%_ = 95% confidence interval), extracted from multivariate linear model fitted in all individuals [metabolite ~ BMI + BMI.score.group + sex + age]. Where an interaction term improved the fit of the model and/or the metabolite level was not associated with measured BMI (*p*>0.05), the measured BMI effect (adjusted for age and sex) is given for each BMI score group separately ($\beta_{BMI\_high}$, $\beta_{BMI\_low}$). In the plots, solid lines denote the predicted univariate within score group relationship between BMI and metabolite with a 95% confidence interval denoted by shading. Metabolites shown in this figure are the eight representative BMI score group associated features that are not included in Figure 5 of the main text.

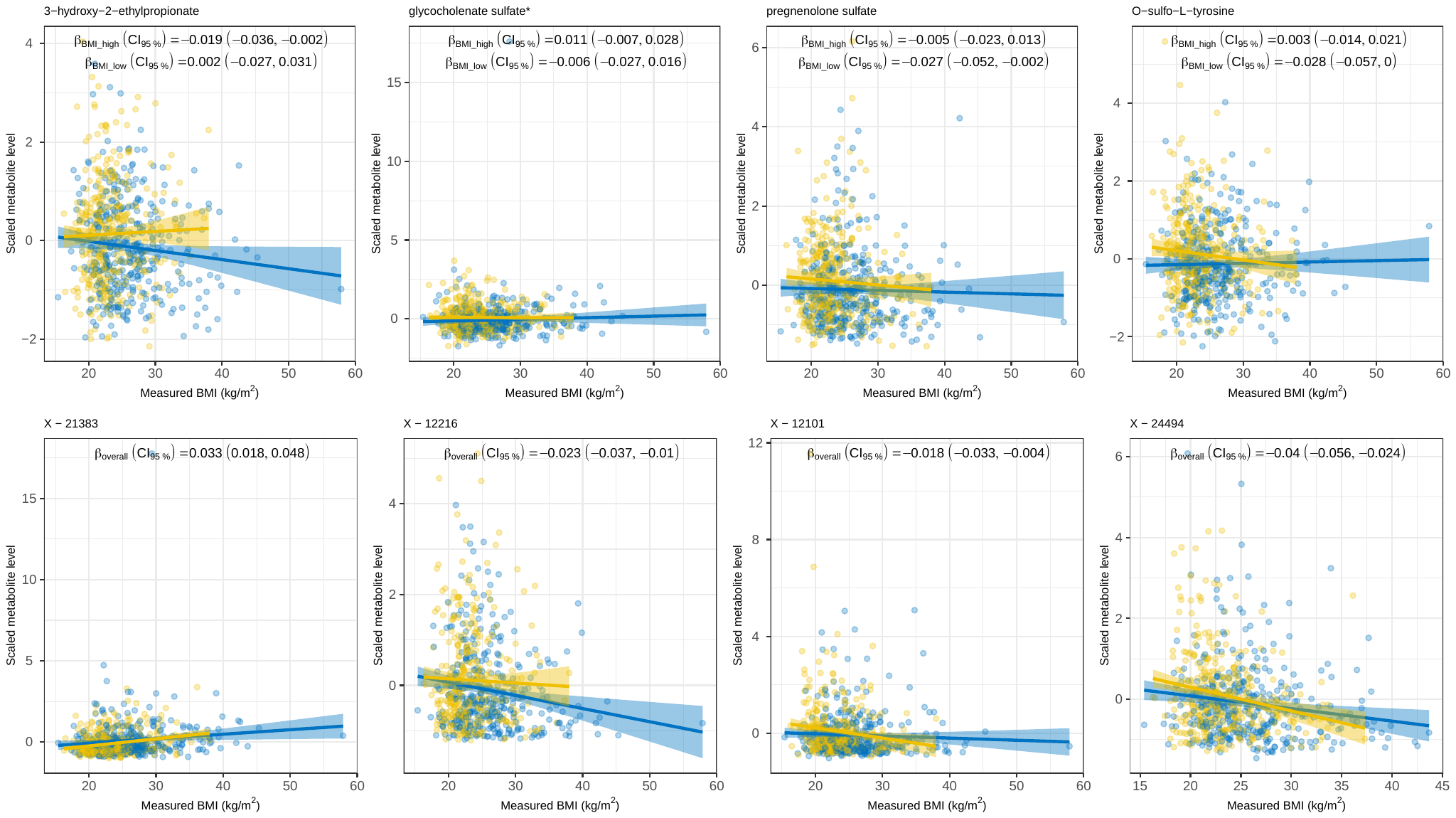
