## Supplementary figures and images for "A multi-variant recall-by-genotype study of the metabolomic signature of body mass index"

### Supporting Information - Figure S3

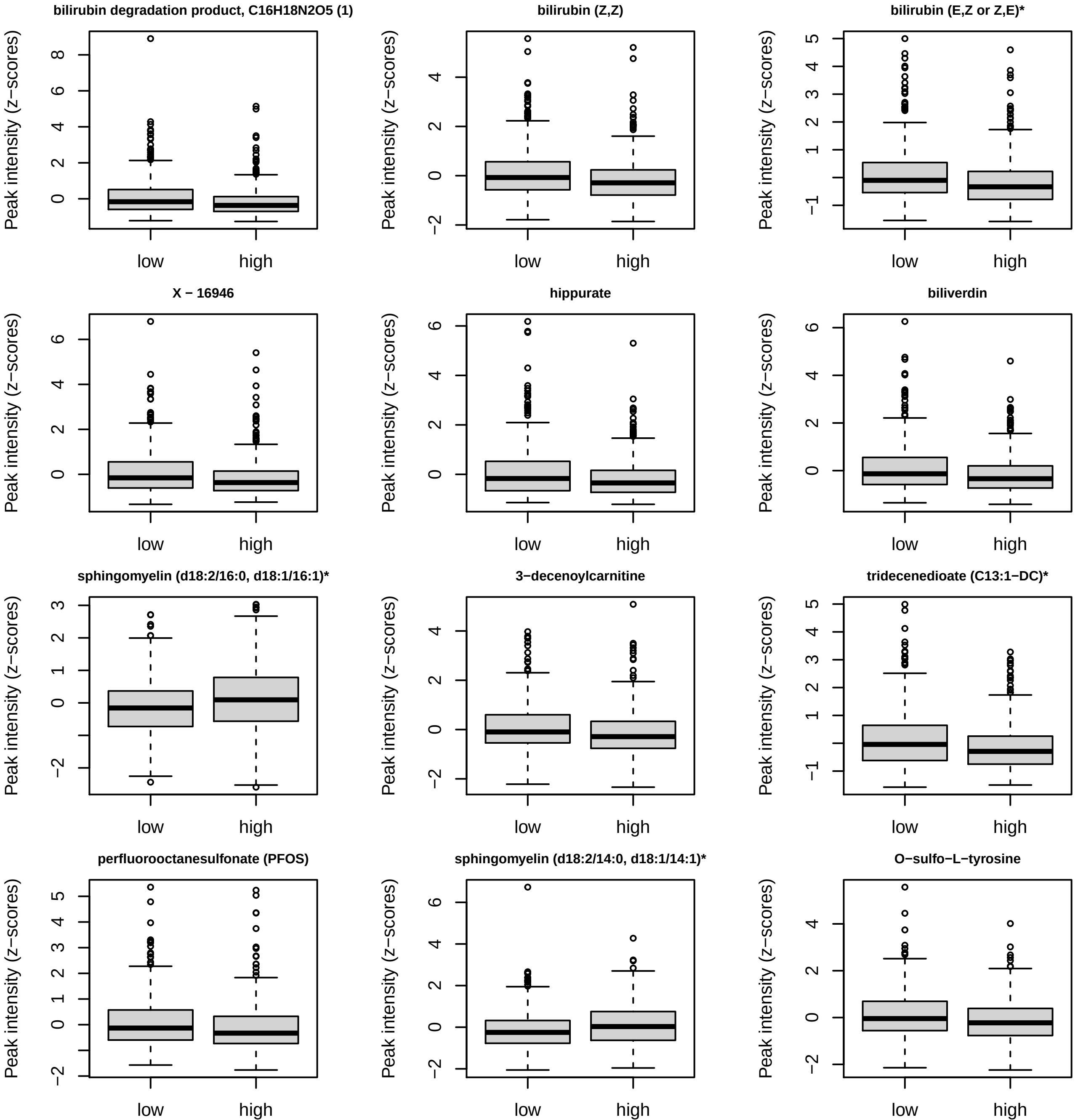

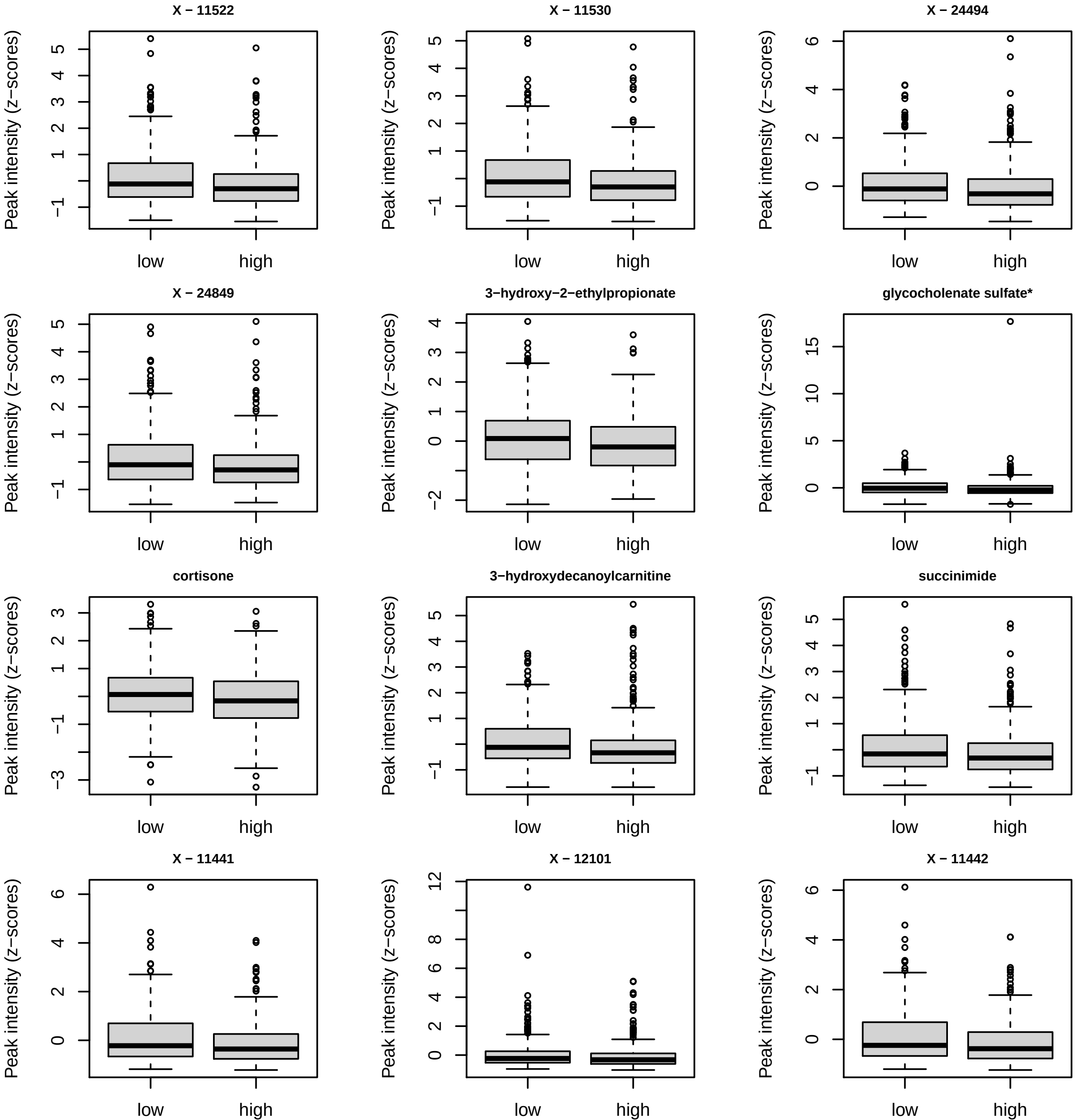

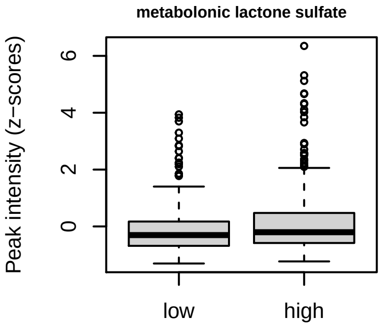

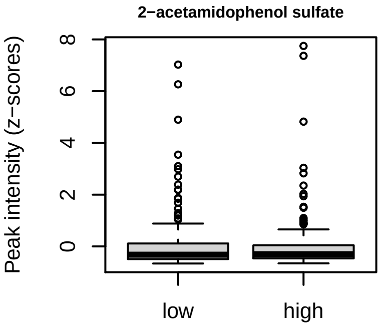
